# Structure tokens sharpen the feature vocabulary of protein language models

**DOI:** 10.64898/2026.05.12.724593

**Authors:** Jacob L Steenwyk

## Abstract

Protein language models predict structure and function from amino acid sequences, but the internal computations that produce these predictions remain opaque. We applied sparse autoencoders to ESM-2 (650M parameters, sequence-only) and ESM-3 (1.4B parameters, multimodal) and found that 78% of learned features converge between the two architectures (permutation null: 14.2%, p < 0.001). These convergent features account for nearly all functional knowledge encoded by the models (functional site AUROC 0.925 versus 0.661 for architecture-unique features). Structure tokens in ESM-3 do not create a new feature vocabulary. Instead, the 15.2% of features most activated by structure tokens are more convergent with sequence-only ESM-2 than structure-invariant features are (r = 0.54 versus 0.45) and carry richer biological annotation (134 versus 29 enriched GO terms). Attention analysis identified a single geometric head (L0H7) as the bottleneck through which structural information enters the network; ablating this head alone changed secondary structure predictions at 40% of residues, while ablating random layer-0 heads altered fewer than 17%. Steering vectors, attribution patching, and sparse feature circuits confirmed that these features sit within the model’s causal pathway. Two architecturally distinct models, trained on different objectives and input modalities, converge on a shared biological vocabulary — and explicit structure tokens sharpen that vocabulary rather than rewriting it.

## 1 Introduction

Protein language models trained on large databases of evolutionary sequences have transformed computational biology. ESM-2, a 650M-parameter sequence-only transformer trained on millions of protein sequences, produces representations that encode secondary structure, binding sites, and long-range contacts without explicit structural supervision [1, 2]. ESM-3, a 1.4B-parameter multimodal model that jointly processes sequence, structure, and function tokens, extends these capabilities to conditional generation of proteins with specified properties [3]. These models are foundational for many structure prediction pipelines, variant-effect prediction, and protein design efforts [1, 4, 5].

However, the computations that give rise to these representations remain poorly understood. When a protein language model assigns high confidence to a particular residue prediction, determining which sequence features, structural regularities, or evolutionary constraints informed the prediction is a challenge. This opacity limits the ability to diagnose model failures, identify the biological knowledge encoded in learned representations, and determine whether models trained on different data modalities converge on the same biological abstractions or arrive at fundamentally different solutions.

Mechanistic interpretability, the effort to reverse-engineer learned computations to a system of human-understandable features and circuits, has made substantial progress in natural language processing [6–8], but its application to protein language models remains in its early stages [9, 10]. One tool for mechanistic interpretability studies is sparse autoencoders (SAEs), which break apart a model’s dense internal representations into a small number of active features, each corresponding to a single interpretable concept [11]. When applied to the internals of a language model, SAEs have identified features corresponding to single syntactic roles, semantic categories, and factual associations [12, 13].

Despite this progress, gaps remain in our understanding of the internals of protein language models. For example, comparing the features learned by architecturally distinct protein language models to determine whether biological knowledge converges across training regimes and model capacities remains poorly understood. Additionally, little is known about multimodal protein language models, leaving open the question of how structure and function tokens reshape the representations that a model learns from sequences alone.

To address these gaps, a comparative SAE framework is introduced that enables systematic feature matching across architecturally distinct protein language models. SAEs were trained on intermediate representations of both ESM-2 (layer 24 of 33) and ESM-3 (layer 33 of 48), using activations extracted from 1.5 million proteins, and evaluated on 12,491 annotated proteins. Complementary mechanistic analyses of attention characterization (layer ablation, attribution patching, and representational similarity) were also applied. Feature quality was assessed through autointerpretability, Gene Ontology enrichment analysis, and functional site detection probing, and revealed that SAEs recovered biologically coherent features. A cross-architecture comparison revealed that 78% of features converge between the two models and that convergent features capture the majority of learned protein biology. Analysis of ESM-3’s multimodal processing revealed that structure tokens reinforced protein biology abstractions learned from sequence alone. Attention analysis across ESM-3 layers revealed a two-stage integration trajectory in which structure-responsive attention heads concentrate in layers 2-9, while representation-level convergence between sequence-only and structure-informed conditions occurs later, across layers 28-38. These findings were validated causally through steering vectors that shift model predictions along specific biological axes, attribution patching that quantifies each layer’s contribution to prediction changes, and interchange interventions that swap representations between proteins with different properties.

## 2 Results

### 2.1 Sparse autoencoders recover biologically meaningful features

Protein language models compress protein biology information into dense, high-dimensional activation vectors that lack direct biological meaning [2, 14]. SAEs enable the decomposition of such representations into individual interpretable units in both natural language models [11, 13] and protein language models [10]. Herein, SAEs were trained on a dataset of 1.5 million UniRef50 proteins at layer 33 of ESM-3 and layer 24 of ESM-2, corresponding to approximately two-thirds network depth [10, 15] (Fig. 1a; Extended Data Fig. 1a-d). The resulting models were evaluated on a holdout set of 12,491 Swiss-Prot proteins with experimentally validated annotations.

**Fig. 1.**
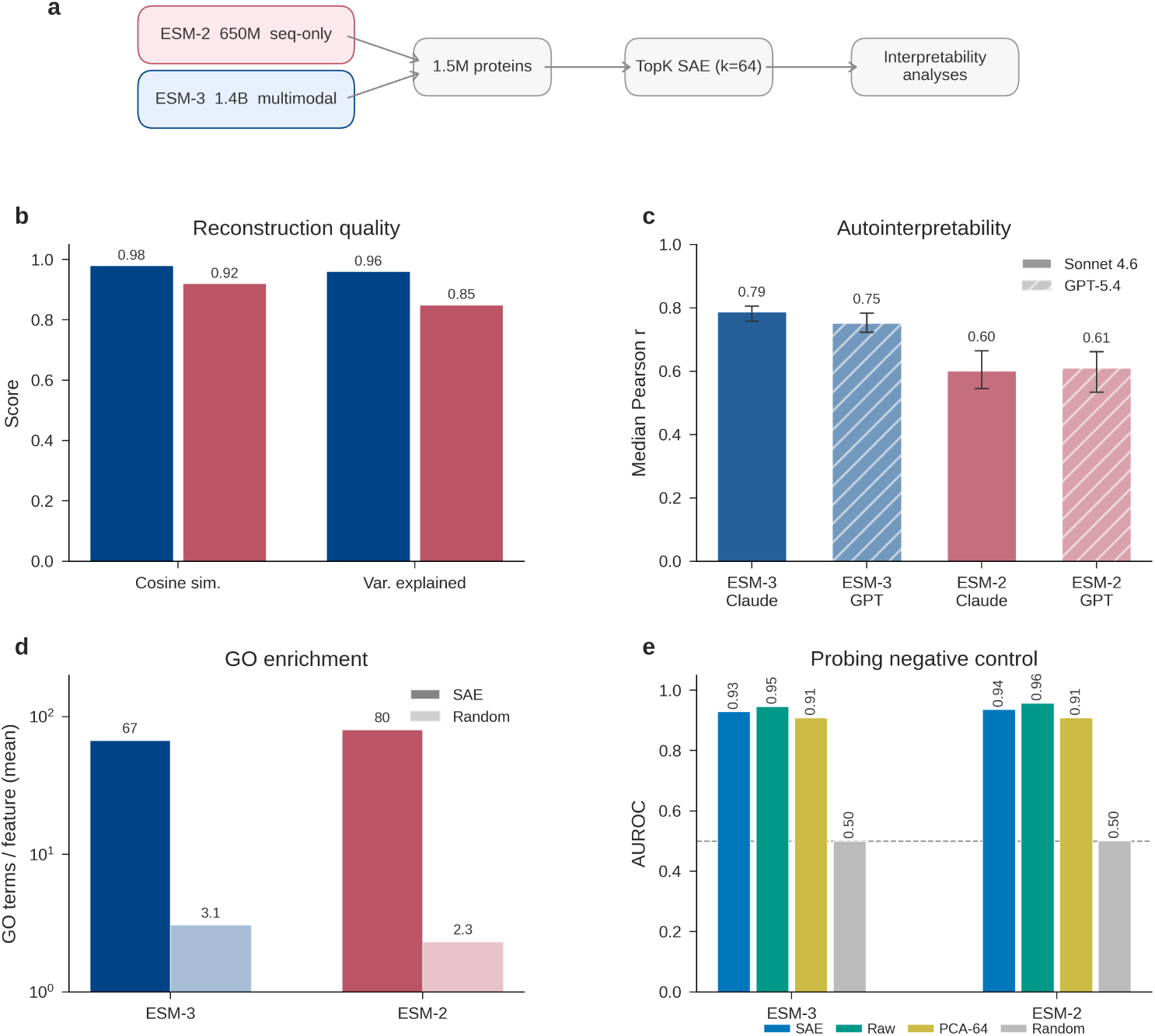
Sparse autoencoders recover biologically meaningful features from protein language models. **a.** SAEs were trained on residual stream activations from 1.5 million UniRef50 proteins decomposing ESM-2 (650M parameters, 33 layers, sequence-only) and ESM-3 (1.4B parameters, 48 layers, multimodal) into sparse features. All evaluations use 12,491 held-out Swiss-Prot proteins unless otherwise noted. **b.** Autointerpretability analysis revealed a median Pearson *r* between LLM-predicted and observed per-residue feature activations on held-out proteins, scored independently by Claude Sonnet 4.6 (solid bars) and GPT-5.4 (hatched bars). ESM-3: *r* = 0.79 (Claude), *r* = 0.75 (GPT); ESM-2: *r* = 0.60 (Claude), *r* = 0.61 (GPT). Error bars show 95% bias-corrected and accelerated bootstrap confidence intervals from protein-level resampling (2,000 iterations). ESM-3 features are substantially more interpretable than ESM-2 features, as judged by both independent judges. **c.** Functional site detection AUROC for linear probes trained on five feature representations: SAE features (ESM-3: 0.93, ESM-2: 0.94), raw full-dimensional activations (0.95, 0.96), PCA with 64 components (0.91, 0.96), shuffled SAE features (0.90, 0.91), and random projections (0.50, 0.50). SAE features match full-dimensional performance while compressing into 64 interpretable dimensions. **d.** Mean number of enriched Gene Ontology terms per feature (BH-FDR *<* 0.05) for SAE features compared to random projection baselines (log *y*-axis). ESM-3: 67 compared to 3.1; ESM-2: 80 compared to 2.3. SAE features are enriched for specific biological functions at 20-30*×* the rate of random baselines. **e.** Reconstruction quality measured by cosine similarity and fraction of variance explained. ESM-3: 0.98 cosine similarity, 96% variance explained (dark blue); ESM-2: 0.92 cosine similarity, 85% variance explained (light red). The sparse bottleneck (64 active features out of 12,288) preserves nearly all representational information.

Reconstruction fidelity was high for both models. The ESM-3 SAEs achieved a cosine similarity of 0.98, indicating how well the SAE reconstructs the original activation vector (Fig. 1b). In addition, SAEs achieved 96% of the activation variance in the evaluation set, which measures how well the reconstruction captures the magnitude and spread of the original activations. ESM-2 SAEs reached 0.92 cosine similarity and 85% variance explained (Fig. 1b; Extended Data Fig. 1f-h). Of the 12,288 learned features in each SAE, 7,459 were active in ESM-3 and 7,533 in ESM-2, leaving 39% and 26% dead features, respectively (Extended Data Fig. 1d). The higher dead-feature rate in ESM-3 likely reflects its larger native dimensionality.

The individual features were then examined for biological meaning using three approaches. The first was autointerpretability analysis, in which a language model generates a natural-language description of each feature from its top-activating residues and a second model scores how well that description predicts held-out activations [10]. The median prediction-activation correlations were higher for ESM-3 features compared to ESM-2 features (r = 0.79 [Sonnet 4.6, Anthropic] and r = 0.75 [GPT-5.4, OpenAI] for ESM-3 and r = 0.60 and r = 0.61 for ESM-2, respectively (Fig. 1c; Extended Data Fig. 2a-d). The two judges strongly agreed on the ranking per-feature (Pearson *r* = 0.91 for the ESM-3 features, *r* = 0.90 for the ESM-2 features), indicating that the interpretability scores reflect the quality of the feature rather than the scoring tendencies specific to the judges. Moreover, the autointerpretability scores reported here are consistent with those obtained by prior SAE analyses of smaller ESM-2 variants (median *r* = 0.72–0.76) [9, 10], validating the approach while extending it to a multimodal architecture and causal validation. The second was Gene Ontology enrichment, in which proteins that strongly activate a feature (top quartile) were examined for functional enrichment. This analysis revealed that 90.3% of active ESM-3 features were enriched for at least one GO term (BH-FDR < 0.05). In contrast, 40.3% of the ESM-2 features had signatures of functional enrichment (Fig. 1d; Extended Data Fig. 2e-h). This gap likely stems from ESM-2 features being more specific, with a median of 5 activation tokens per feature, compared to 5,697 for ESM-3, making enrichment testing underpowered for most ESM-2 features.

The third approach examined whether SAE features preserve sufficient information for biological prediction. Linear probes trained on SAE activations for functional site detection achieved AUROC 0.93, matching the performance of probes on raw, full-dimensional activations (0.94) (random baseline: 0.50; Fig. 1e). A PCA baseline using the same number of components (64) achieved comparable AUROC (0.91), confirming that the SAE does not sacrifice predictive power for interpretability (Extended Data Fig. 3a-b).

### 2.2 Feature representations converge across architectures

ESM-2 and ESM-3 differ in parameter count (650M and 1.4B, respectively), architecture, input modalities, and training data. ESM-2 uses masked language modeling in sequences alone, while ESM-3 is trained with a generative objective in sequence, structure, and function [1, 3]. If the constraints of protein biology are strong enough to overcome these architectural and training differences, the two models should learn similar internal features.

To test this, the Pearson correlation between the protein-level activation profile of the 7,459 active ESM-3 features and the 7,533 active ESM-2 features across the shared evaluation proteins was computed, and the best match was retained. The median best-match correlation was r = 0.456, and 78% of ESM-3 features found an ESM-2 partner above r > 0.3 (Fig. 2a). A permutation null constructed by shuffling protein labels yielded a median of r = 0.165, with only 14.2% of features crossing the r > 0.3 threshold (p < 0.001, Cohen’s h = 1.39; Fig. 2b). Results were robust to homology-aware splits, with probing AUROC holding at 0.923 across five 50% sequence identity folds (Extended Data Fig. 3d-e). Because many features are zero for most proteins and shared zeros could inflate Pearson correlations, the analysis was repeated using Spearman’s rank correlation, revealing negligible inflation (+0.005; from 0.456 to approximately 0.461).

**Fig. 2.**
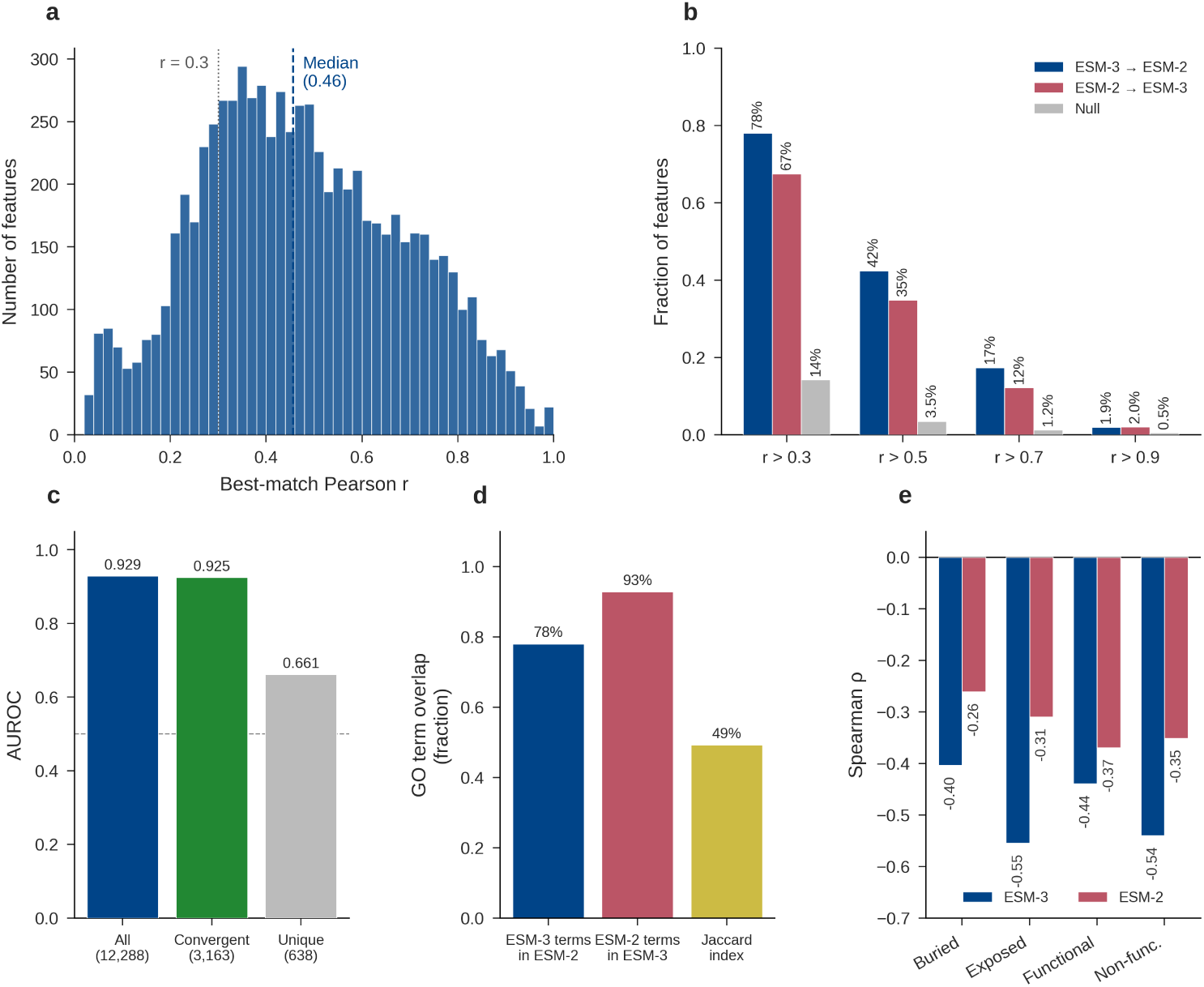
Feature representations converge across architectures. **a.** Distribution of best-match Pearson *r* between each ESM-3 feature and all ESM-2 features, computed over protein-level activation profiles. Dashed lines mark the median (*r* = 0.46) and the *r* = 0.3 convergence threshold. The majority of ESM-3 features find a correlated partner in ESM-2 despite the two models differing in architecture, scale, and input modality. **b.** Fraction of features exceeding convergence thresholds (*r >* 0.3, *>* 0.5, *>* 0.7, *>* 0.9) for ESM-3 to ESM-2 matching (dark blue), ESM-2 to ESM-3 matching (light red), and a permutation null with shuffled protein labels (grey). At *r >* 0.3, 78% of features converge compared to 14% in the null (binomial *p <* 0.001, Cohen’s *h* = 1.39). Convergence exceeds chance at every threshold tested. **c.** Functional site detection AUROC stratified by convergence. Convergent features (*r >* 0.5, *n* = 3,163; AUROC 0.925) retain 99.6% of full-feature-set performance (*n* = 12,288; AUROC 0.929), whereas architecture-unique features (*r <* 0.2, *n* = 638; AUROC 0.661) carry less biological signal. The features both models agree on are the ones that encode biology. **d.** Absolute Spearman *|ρ|* between SAE activation magnitude and masked language model entropy for buried, exposed, functional, and non-functional residue classes in ESM-3 (dark blue) and ESM-2 (light red). Both models encode evolutionary conservation across residue categories. **e.** GO term overlap between matched ESM-3 and ESM-2 feature pairs. ESM-3 terms recovered in ESM-2: 78%. ESM-2 terms recovered in ESM-3: 93%. Mean Jaccard index: 0.49. Matched features share specific biological annotations.

Finding the most similar feature in each respective model was influenced by the directionality of the comparison. In the ESM-3 to ESM-2 direction, 78% of features were convergent (r > 0.3). In the reverse direction (ESM-2 to ESM-3), 67% were convergent (Fig. 2b). This asymmetry is consistent with ESM-3’s larger model capacity and multimodal training. More specifically, ESM-3 represents everything ESM-2 learns, plus additional features (likely structure- and function-track representations) not represented in ESM-2.

To determine whether convergent features are more biologically informative, the ESM-3 feature set was split by match quality, and separate functional site probes were trained. Features with high cross-model correlation (r > 0.5; n = 3,163) achieved AUROC 0.925, which corresponds to 99.6% of the complete feature set (AUROC = 0.929). Architecture-unique features (r < 0.2; n = 638) achieved an AUROC of 0.661 (Fig. 2c). The features that both models agree on are, overwhelmingly, the features that matter for biology. GO term overlap analysis reinforced this conclusion. The Jaccard index between the GO term sets of matched ESM-3 and ESM-2 features was 0.492, with 78% of ESM-3 feature GO terms appearing in the matched ESM-2 feature and 93% of ESM-2 GO terms appearing in the matched ESM-3 feature (Fig. 2d).

A further analysis examined whether convergence tracks with evolutionary constraint. SAE activation magnitudes correlated negatively with positional entropy across all residue categories (*ρ* = −0.26 to −0.55), indicating that both models activate features more strongly at evolutionarily conserved positions (Fig. 2e). ESM-3 showed consistently stronger correlations than ESM-2, with the largest gap at exposed (*ρ* = −0.55 versus −0.31) and non-functional (*ρ* = −0.54 versus −0.35) positions. The two models converged most closely at functional sites (Δ*ρ* = 0.07; *ρ* = −0.44 versus −0.37), where sequence conservation provides a strong and direct biological signal, and diverged most at non-functional sites (Δ*ρ* = 0.19), where conservation signals are weaker and more distributed. This pattern suggests that both architectures converge when the biological signal is strong, while ESM-3’s advantage is most apparent at positions where the signal is subtle — though the current analysis does not distinguish whether the gap reflects ESM-3’s larger parameter count, its access to structure tokens, or both.

Feature breadth predicted convergence strength. Generalist features activating on more than 10,000 tokens converged strongly (median r = 0.48), while specialist features activating on fewer than 100 tokens did not (r = 0.30; Mann-Whitney p < 0.001). Broad biological properties, such as secondary structure, solvent accessibility, and charge, are recovered by both architectures. For example, proteins with WD40 repeats and chromatin organization proteins converged between the two architectures (r = 0.81). In contrast, specialist features, such as bacterial inner membrane proteins involved in cell surface biogenesis (r = 0.03) and glycosyltransferases involved in glycan biosynthesis (r = 0.07), did not. This gradient from universal to architecture-specific features suggests that convergence is driven by the frequency and consistency of biological signals in the training data, not by any structural similarity between ESM-2 and ESM-3 (Extended Data Fig. 4).

### 2.3 Multimodal integration has a two-stage trajectory with early attention and late convergence

ESM-3 can accept structure tokens (representations of 3-dimensional coordinates) along with the amino acid sequence [3]. Whether and how this structural information reshapes the model’s internal representations remains unclear. To examine this, ESM-3 was run under sequence-only (S) and sequence-plus-structure (S+St) conditions on 991 proteins with predicted structures of AlphaFold [16]. Subsequently, the centered kernel alignment (CKA), which measures how similar two sets of representations are in terms of their geometric structure (where values from zero to one and high values indicate greater similarity), was examined between the two activation sets at 12 layers spanning the full network (Fig. 3a).

**Fig. 3.**
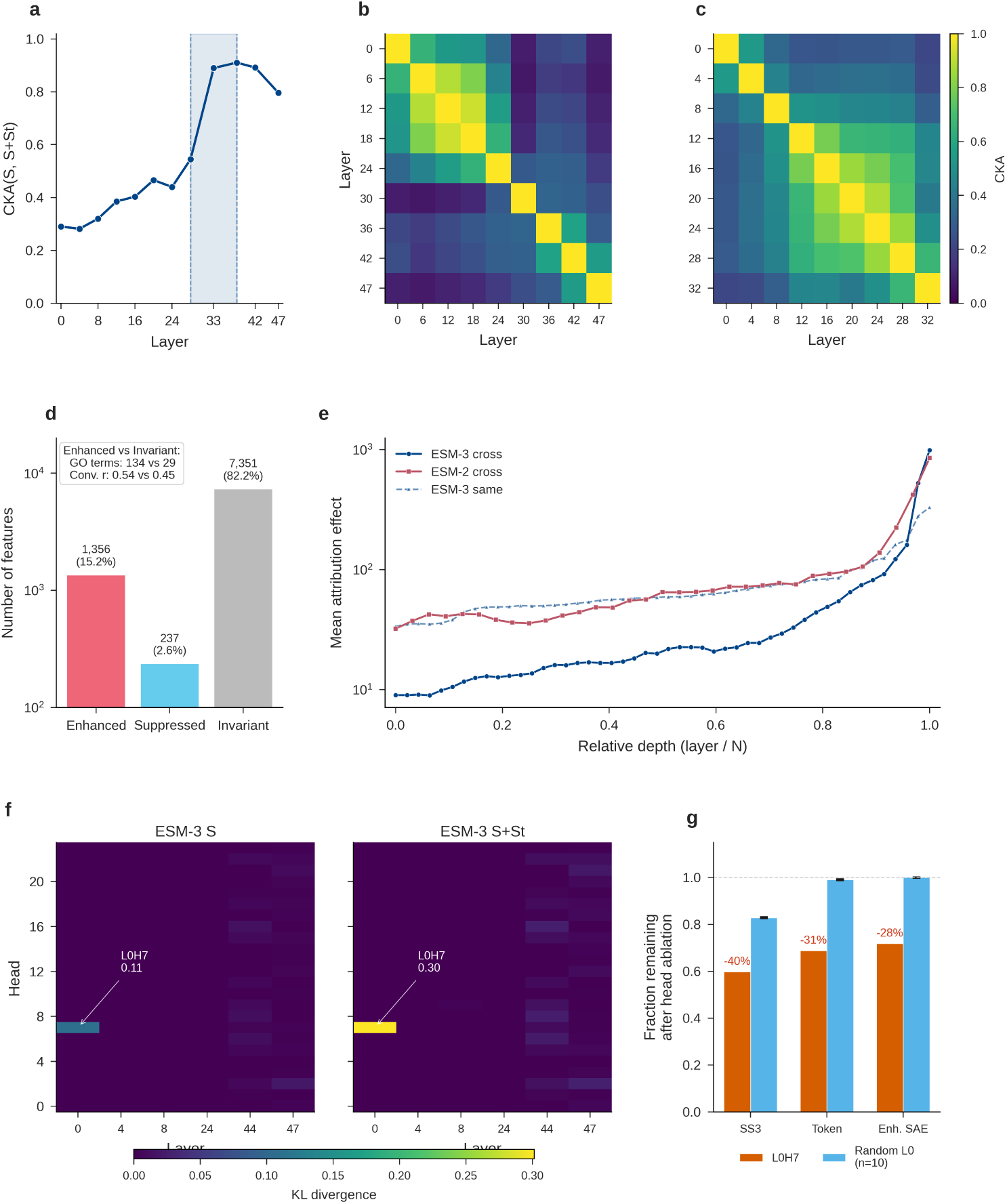
Structure tokens reshape a minority of features through early attention and late representational convergence. **a.** CKA between sequence-only (S) and sequence-plus-structure (S+St) activations across ESM-3 layers (991 proteins with AlphaFold structures). CKA rises from 0.29 at layer 0 to 0.91 at layer 38, then declines to 0.80 at layer 47. Shaded region: integration zone (layers 28-38), where structure information is reconciled into a unified representation. **b.** Cross-modal feature categories among 8,944 active features at layer 33: structure-enhanced (1,356, 15.2%), structure-suppressed (237, 2.6%), and invariant (7,351, 82.2%). Log *y*-axis. Inset: enhanced features carry more GO terms (134 compared to 29) and higher cross-architecture convergence (*r* = 0.54 compared to 0.45) than invariant features. Structure tokens reshape a minority of features, but those features are disproportionately biologically informative. **c.** Attribution patching effect by relative network depth for 200 helix-versus-sheet protein pairs. ESM-3 cross-protein (blue, solid), ESM-2 cross-protein (red, solid), ESM-3 same-protein control (blue, dashed). Log *y*-axis. Both models concentrate structural discrimination in their final layers. **d.** Head ablation KL divergence heatmaps (layer *×* head) from zeroing individual attention heads under S (left) and S+St (right). L0H7 KL increases from 0.11 (S) to 0.30 (S+St), identifying a single geometric attention head as the primary bottleneck for structural information. **e.** Functional impact of L0H7 ablation under S+St. Fraction remaining after ablating L0H7 (orange) compared to 10 random layer-0 heads (blue, mean *±* s.d.). L0H7: *−*40% SS3 accuracy, *−*31% token predictions, *−*28% enhanced SAE features. Random heads have a negligible impact.

Structure tokens produce representations different from those computed from sequence alone. At layer 0, CKA(S, S+St) is 0.29, reflecting the distinct embedding spaces that sequence and structure tokens occupy. The two representation streams remain largely dissimilar through the first half of the network (CKA < 0.47 through layer 24), then converge; CKA reaches 0.89 at layer 33, peaks at 0.91 at layer 38, and drops back to 0.80 at layer 47 (Fig. 3a). The integration zone (layers 28-38) may correspond to the point at which the model integrates the two input modalities into a unified representation. The decline in layer 47 suggests that the information derived from the structure is deployed differentially at the output head (Extended Data Fig. 5). Within-model CKA independently supports this boundary: ESM-3 undergoes an abrupt representational phase transition between layers 24 and 30 (CKA drops to 0.08), whereas ESM-2 evolves representations more gradually (Fig. 3b,c).

To determine how structure tokens alter individual SAE features, each of the 8,944 active features in layer 33 was classified by comparing its activation profile in 11,704 proteins (a subset of the 12,491 evaluation proteins that had structures predicted by AlphaFold [16]) run under both S and S+St conditions (Wilcoxon signed-rank test with BH correction). Of these, 1,356 (15.2%) were structure-enhanced (higher activation with structure tokens), 237 (2.6%) were structure-suppressed (lower activation with structure tokens), and 7,351 (82.2%) were invariant (Fig. 3d; Extended Data Fig. 6). The enhanced features were more convergent with sequence-only ESM-2 than the invariant features (median cross-architecture r = 0.54 compared to r = 0.45; Mann-Whitney p < 0.001), and they carried richer biological functions (median 134 enriched GO terms compared to 29 for invariant features; p < 0.001; Fig. 3d inset). These findings indicate that the features that structure tokens strengthen are the same features that ESM-2 learns to activate through implicit structural inference, suggesting that structure tokens, which represent a new modality, do not introduce a new feature vocabulary that sequence alone does not capture.

To further examine this, attribution patching was conducted. Attribution patching quantifies each layer’s causal contribution to a prediction by estimating how much the output would change if the activations of that layer were swapped between two proteins. Attribution patching was applied to 200 protein pairs in which one protein was helix-dominant and the other sheet-dominant. Both models concentrated structural discrimination in their final layers (ESM-3: effect 9.0 at layer 0 to 991 at layer 47; ESM-2: 32 to 853; Fig. 3e). ESM-3’s 3.6-fold lower early-layer effect reflects the structure embedding itself because explicit 3-dimensional tokens reduce the need for early layers to infer secondary structure from sequence context. Running attribution patching on the same protein under S compared to S+St conditions produced a similar late-layer peak (Fig. 3e, light blue; Extended Data Fig. 3c), supporting that the cross-protein result reflects structure processing rather than sequence differences between paired proteins.

A complementary analysis examined which attention heads mediate structure integration. By computing the Jensen-Shannon divergence between each head’s attention pattern under S and S+St conditions across 494 proteins, 28 structure-responsive heads were identified (JSD > 0.1), all concentrated in layers 2-9 (mean JSD 0.09-0.15; Extended Data Fig. 7a-c). Attention patterns in the final third of the network (layers 32-47) were identical regardless of whether structure tokens were provided. The contact-predicting heads, by contrast, reside in layers 44-47 (top head: L45H11, precision 0.60 at L-cutoff; Extended Data Fig. 8a-c). The heads that respond to structural input and the heads that predict 3D contacts are therefore distinct populations operating at opposite ends of the network, with early heads reading structure in and late heads reading structure out.

Head ablation causally confirmed this architecture. When individual heads were zeroed, and the resulting KL divergence from unperturbed output was measured across 100 proteins under both S and S+St conditions, head L0H7 emerged as the single most critical head (Fig. 3f). Its importance nearly tripled when structure tokens were provided (KL 0.11 under S, 0.30 under S+St), consistent with its role as the primary head for ingesting 3-dimensional coordinate information. Ablating L0H7 alone under S+St had broad functional consequences (Fig. 3g). Specifically, 40% of residues received a different secondary structure prediction (helix, sheet, or coil), 31% of amino acid predictions changed, and structure-enhanced SAE features lost 28% of their mean activation. A comparison against 10 randomly selected layer-0 heads confirmed that L0H7’s importance is specific; ablating random heads preserved 83% of secondary structure predictions, 99% of amino acid predictions, and 99.9% of SAE feature activations.

### 2.4 SAE features causally mediate model predictions

To test whether the features identified by the trained SAEs causally mediate the model behavior, three intervention classes were implemented. First, the model was steered along a specific biological direction. A steering vector was constructed by averaging the decoder directions of the 20 most structure-enhanced SAE features. During inference on 500 proteins, this vector was injected into the model’s intermediate representations at varying strengths (*α* = -50 to +50), amplifying or suppressing the targeted features (Fig. 4a). The result is dose-dependent, with target-feature activation scaling with steering strength (r = 0.87). Moreover, the effect was symmetric, with steering in either direction producing proportional changes. At the strongest perturbation (*α* = +50), 4.0% of the predicted amino acid tokens changed identity, confirming that these features influence what the model predicts. Random, shuffled, and orthogonal directions produced lower KL at matched steering strength (Fig. 4b), supporting that the effect is specific to the learned feature direction.

**Fig. 4.**
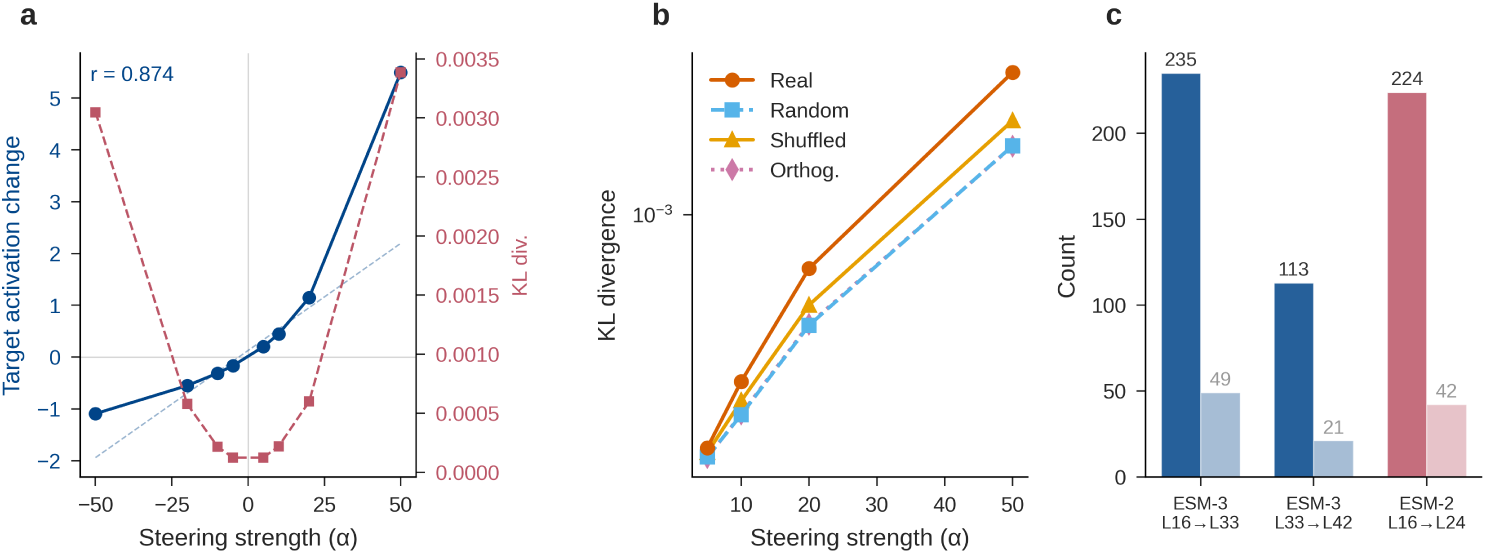
SAE features causally mediate model predictions. **a.** Steering dose-response. A steering vector constructed from the mean decoder direction of the 20 most structure-enhanced SAE features is added to the ESM-3 residual stream at layer 33 with scaling factor *α* from *−*50 to +50. Left axis (blue): target feature activation change (*r* = 0.87 with *α*). Right axis (red): KL divergence from unsteered predictions. Feature activation scales linearly and symmetrically with steering strength, confirming dose-dependent causal control. **b.** Specificity controls. KL divergence across steering strengths for real SAE decoder directions (solid orange), random directions (dashed blue), shuffled decoder columns (solid yellow), and orthogonalized directions (dotted pink). Log *y*-axis. Real directions produce larger effects than all controls at every strength tested, confirming that the steering effect is specific to the learned feature direction rather than a generic response to residual stream perturbation. **c.** Sparse feature circuits. Number of significant causal connections (dark bars) and downstream features with at least one upstream driver (light bars) per layer pair, determined by single-feature ablation with Bonferroni-corrected *t*-tests. ESM-3 L16*→*L33: 235 connections driving 49 of 50 features; L33*→*L42: 113 connections driving 21 of 50. ESM-2 L16*→*L24: 224 connections driving 42 of 50. Features form causal processing chains across layers, with decreasing connection density from early-to-mid to mid-to-late pairs.

Second, causal wiring between features was mapped at different layers. If SAE features constitute a processing sequence in which each step builds on the last, then ablating an upstream feature should alter the activation of downstream features that depend on it. This was tested by systematically removing individual features in an earlier layer and measuring the effect on features in a later layer, retaining only connections that were significant by Bonferroni-corrected *t*-tests and exceeded an effect size threshold of two standard deviations above the null median (Fig. 4c). In ESM-3, 235 causal connections linked layer 16 features to layer 33 features, and 113 causal connections linked layer 33 to layer 42. ESM-2 showed a similar density (224 connections from layer 16 to layer 24). Of the 50 downstream features tested at each layer pair, 49, 21, and 42 had at least one significant upstream driver, respectively. This finding indicates that nearly all mid-layer features depend on earlier computation, while fewer late-layer features require active mid-layer input. The decreasing connection density from early-to-mid (235) to mid-to-late (113) is consistent with the progressive representational convergence observed in the CKA analysis (Extended Data Fig. 8d-e; Extended Data Fig. 9).

Third, the features were grounded in specific proteins with well-characterized biology. Three proteins — a metalloprotease and two cytochromes — were selected because their functional sites are densely annotated (Extended Data Fig. 10). For each protein, per-residue SAE feature activations were overlaid on experimentally validated functional site positions from UniProt. In serralysin, structure-enhanced features activated at the zinc-binding site and calcium-coordinating residues required for catalytic activity (Extended Data Fig. 10a). In the two multi-heme cytochromes, a shared set of features (F435, F16) tracked the regularly spaced heme-binding CXXCH motifs, indicating that the SAE has learned a general heme-coordination feature that generalizes across protein families (Extended Data Fig. 10b,c). These case studies confirm that the features flagged as structure-enhanced by the statistical analysis localize to the specific residues where structure matters most for function.

Taken together, the steering, circuit, and case-study analyses provide evidence that the SAE features identified herein are chained together to propagate information across layers. The SAE features also respond predictably to targeted perturbation, and they localize to residues where structure and function intersect.

## 3 Discussion

Sparse autoencoders decompose the dense representations of protein language models into features that are individually interpretable and collectively account for the biological knowledge these models encode. By applying this decomposition to two architecturally distinct protein language models (ESM-2, 650M parameters, sequence-only, and ESM-3, 1.4B parameters, multimodal) and validating the resulting features through Gene Ontology enrichment, functional site detection, and causal interventions, a comparative framework is established and distinguishes shared biological abstractions from those that are model-specific. The features that emerge capture independently recovered and shared principles of protein biology.

The degree of feature convergence between these two models suggests that sequence alone achieves a rich representation of protein biology. Specifically, the observation that 78% of learned features are shared across a 650M-parameter sequence-only transformer and a 1.4B-parameter multimodal model (permutation null of 14.2%) suggests that protein language models converge on a shared representational vocabulary regardless of architecture or input modality (Fig. 2a-b). For example, convergent features achieve functional site detection AUROC of 0.925 compared with 0.661 for architecture-unique features (Fig. 2c). Broadly activating generalist features converge, whereas specialist features do not. A similar pattern has been reported in natural language models, where SAEs recover overlapping feature sets across model families [13], suggesting that convergent feature learning reflects a shared property of transformers trained on data with rich combinatorial structure.

An unanticipated finding concerns the relationship between multimodal inputs and feature convergence. Features that have different activation patterns with structure in ESM-3 are more convergent with sequence-only ESM-2 (r = 0.54 compared to 0.45 for structure-invariant features) and more enriched for Gene Ontology terms (134 compared to 29 terms) (Fig. 3d). At face value, one might expect features that respond to an input modality absent from ESM-2 to diverge from ESM-2’s representations; however, findings herein reveal convergence. This may be explained by ESM-2 having to infer structural regularities from sequence context alone, and it does so with sufficient accuracy that its representations already encode the same information that ESM-3 receives explicitly through structure tokens. This suggests that structure tokens do not introduce new vocabulary (or lexicon). Instead, structure tokens sharpen representations that sequence context has already approximated. In practice, this implies that multimodal training may improve the precision of structural encoding without expanding the scope of biological knowledge captured by the model’s features.

Beyond feature identity, the causal analyses constrain where and how these features operate within each model. CKA can be sensitive to a small number of high-variance principal components [17], but the integration-zone boundary is independently supported by the abrupt within-model phase transition in ESM-3 (Fig. 3b,c) and by attribution patching (Fig. 3e), which identifies the same late-layer concentration through a causal rather than geometric method. Both models concentrate prediction formation in the final layer (ESM-2 layer 32, ESM-3 layer 47) as revealed by logit attribution and attribution patching (Fig. 3e; Extended Data Fig. 5). The attention heads that respond to structure tokens are distinct from those that predict residue contacts [14], indicating functional specialization within the attention mechanism. These observations constrain where targeted interventions, such as feature ablation or steering (Fig. 4a,b), are most likely to alter model behavior, and can inform applications such as model editing and guided protein design.

Ablating a single attention head (L0H7) altered 40% of secondary structure predictions and reduced structure-enhanced SAE feature activations by 28% (Fig. 3f,g). In contrast, ten randomly chosen layer-0 heads altered fewer than 17% of predictions and left SAE features intact (99.9% preserved). One head out of 1,152 carries disproportionate responsibility for the input modality. This concentration is unusual; modality-specific computation in transformers is typically distributed across head populations. Whether narrow bottleneck heads arise generally in multimodal architectures or reflect ESM-3’s training regime, in which structure tokens are a minority modality added to a primarily sequence-based objective, remains untested. L0H7’s existence does imply that targeted single-head interventions can selectively diminish modality-specific reasoning from a multimodal model without retraining. This observation is directly relevant to model editing and controlled comparisons between sequence-only and multimodal inference.

Several limitations qualify these conclusions. The SAE analysis examined single layers (layer 24 of 33 for ESM-2, layer 33 of 48 for ESM-3), and multi-layer analyses would reveal how features change across depth. Moreover, wet-lab validation of the biological roles assigned to individual features remains untested and is likely to be of interest. The present comparison spans two models, and extending the framework to other protein-language model families would reveal whether any insights are more broadly applicable. In addition, steering vectors produce dose-responsive shifts in model predictions, but the magnitudes are small, leaving open the question of whether stronger interventions would reveal qualitatively different behavior or scale the observed effects.

A methodological contribution of this work is a comparative SAE framework for distinguishing universal from architecture-specific biological representations across protein language models. The framework consists of three steps: training matched SAEs on intermediate representations of two or more models, matching features across models by protein-level activation profiles, and stratifying results by match quality to separate shared from model-specific signal. This protocol generalizes to other protein language model families such as ProtGPT2 [18], SaProt [19], and ProGen2 [5], and provides a systematic means of identifying which aspects of biological knowledge are universal to the protein modeling task and which are contingent on particular training decisions.

## Methods

### Protein language models

Two protein language models that differ in scale, architecture, and input modalities were analysed. ESM-2 (*facebook/esm*2_*t*33_650*M* _*UR*50*D*) is a 650M-parameter, 33-layer, sequence-only transformer with *d_m_odel* = 1280 and 20 attention heads per layer [1]. ESM-2 was loaded through the HuggingFace transformers library, which exposes attention weights natively via the output_attentions flag. ESM-3 (*esm*3_*sm*_*open*_*v*1) is a 1.4B-parameter, 48-layer multimodal transformer with *d_m_odel* = 1536 and 24 attention heads per layer [3]. ESM-3 was loaded through the EvolutionaryScale esm package, v3.2.1, and all ESM-3 computations were run under torch.autocast(“cuda”, dtype=torch.bfloat16) to match the model’s native precision.

ESM-3 accepts three input modalities [3]. Sequence tokens are standard amino acid indices. Structure tokens are produced by a vector-quantized variational autoencoder (VQ-VAE) that encodes 3D backbone coordinates into a 4,096-entry code, yielding one discrete structural token per residue. Function tokens encode InterPro domain annotations through a pipeline that converts InterPro identifiers [20] to TF-IDF vectors and then applies locality-sensitive hashing (LSH) to produce an integer tensor (L, 8) per protein, where L is the sequence length and the vocabulary size is 260. The three types of token are embedded independently and summed at the model input, after which they share a single transformer trunk with rotary positional embeddings (RoPE) [21].

### Training dataset

Sparse autoencoder training used activations extracted from 1.5 million proteins sampled from UniRef50 (download date: February 23, 2026) [22]. The sequences were filtered to 50-1,022 amino acids, containing only 20 standard residues, and then uniformly sampled from the remaining pool. Each protein was passed through the target model once to extract residual stream activations in the chosen layer. For ESM-3, the chosen layer was layer 33 of 48 total. For ESM-2, the chosen layer was layer 24 of 33 total. The resulting per-residue vectors were pooled across all proteins to form the SAE training set.

### Evaluation dataset

A held-out evaluation set of 12,491 proteins was drawn from Swiss-Prot (reviewed entries only) [23]. To prevent information leakage, evaluation sequences were clustered against the 1.5M training set using MMseqs2, v6e3c640418f9cde4b246f430322e17be7b96a3b9 [24], at 50% sequence identity, and any evaluation protein that fell into a training cluster was removed. The remaining proteins were sampled with taxonomic stratification to preserve approximate kingdom-level proportions of the full Swiss-Prot distribution. Gene Ontology (GO) terms [25], functional site annotations (active sites, binding sites, metal binding), and Pfam domain assignments [26] were retrieved from the UniProt REST API [22, 27].

### Sparse autoencoder architecture

TopK sparse autoencoders [11, 13, 28] were used, which enforce exact sparsity by retaining only the *k* largest encoder activations and zeroing the rest. The encoder is a linear projection from *d*_*model* to *d*_*sae* followed by TopK selection and a ReLU nonlinearity applied to the surviving entries; the decoder is a linear projection from *d*_*sae* back to *d*_*model*. Both encoder and decoder have learned bias terms, and a pre-bias parameter is subtracted from the input before encoding and added back after decoding. SAEs were implemented in PyTorch, v2.6.0+cu124 [29], and trained using the Adam optimiser. The hyperparameters were set to *k* = 64 and expansion factor 8 for all SAEs, yielding *d*_*sae* = 12,288 for ESM-3 (1536 × 8) and *d*_*sae* = 10,240 for ESM-2 (1280 × 8). SAEs were trained on layer 33 of ESM-3 (of 48 total), with activations extracted using the EvolutionaryScale esm package, v3.2.1 [3], and layer 24 of ESM-2 (of 33 total), with activations extracted using HuggingFace transformers, v4.48.1 [30]. These correspond to the approximate two-thirds depth point in each model, where prior work has identified the richest functional information [10, 15].

### Sparse autoencoder training

SAEs were trained with the Adam optimizer at a learning rate of 3 × 10^−4^, with linear warmup over the first 1,000 steps and no weight decay. Batch size was set to 4,096 residue vectors per step. Decoder columns were renormalized to unit *ℓ*_2_ norm after each gradient step. Dead features, defined as dictionary elements that never activated across a random sample of 10 batches, were monitored and resampled every 500 steps by reinitializing their encoder and decoder weights. Training ran for 10 epochs over the respective activation sets, corresponding to approximately 23,000 steps for ESM-3 and 46,000 steps for ESM-2. A 10% held-out validation split was used for early stopping based on reconstruction loss.

### Autointerpretability

Individual SAE features were assessed for their correspondence with human-understandable biological concepts using the an established autointerpretability protocol [9]. For each of 300 randomly sampled features per model, the top-activating residues were identified in a *description set* of 9,000 proteins (drawn from the evaluation set), and Claude Sonnet 4.6 (Anthropic), accessed via the Anthropic Python SDK, v0.84.0, was prompted to generate a natural-language description of the activation pattern. The description was scored by two independent language models — Claude Sonnet 4.6 and GPT-5.4 (OpenAI, accessed via the OpenAI Python SDK, v2.26.0) — each given the description and a *validation set* of 3,491 held-out proteins and asked to predict, for each residue, how strongly the feature would activate on a 0–1 scale. The Pearson correlation between predicted and actual activation values was calculated using SciPy v1.13.1 [31] for all residues in the validation set, with missing predictions set to 0.5 (a conservative default that biases the metric towards zero correlation). The description and validation sets were disjoint to prevent memorization.

### Gene Ontology enrichment analysis

For each active SAE feature, the study set was defined as the set of proteins whose maximum per-protein activation for that feature fell in the top quartile. The population was all 11,288 evaluation proteins with at least one GO annotation. Over-representation analysis was performed using the gokit library, v0.1.3 https://github.com/JLSteenwyk/gokit, applying the Benjamini-Hochberg false discovery rate correction (method=“fdr_bh”) [32]. A feature was considered enriched for a GO term if the adjusted p-value was below 0.05. Odds ratios were computed with a floor of 10^−6^ in the denominator to avoid division by zero for terms absent from the background. GO annotations were retrieved from UniProt REST API [22, 27].

### Cross-architecture feature convergence

To test whether ESM-2 and ESM-3 learn overlapping feature vocabularies, Pearson correlations were computed using NumPy, v2.1.0 [33], between the protein-level maximum activation profiles of all active ESM-3 features and all active ESM-2 features across the 12,491 shared evaluation proteins. For each ESM-3 feature, the best-match *r* with any ESM-2 feature was retained. A feature was classified as convergent if *r >* 0.3. Spearman rank correlations were computed using SciPy, v1.13.1 [31], to confirm that results were not driven by outliers. A permutation null distribution was constructed by shuffling protein labels 10 times and recomputing best-match *r* values. The observed convergence rate was compared against the resulting null using a binomial test (SciPy stats.binom_test). To rule out the possibility that shared zeros inflated Pearson correlations (both features inactive on the same proteins), the analysis was repeated after excluding all protein pairs where both features were zero; the resulting correlations differed by only 0.005 on average.

### Cross-modal feature analysis

Of the 12,491 evaluation proteins, 11,704 had AlphaFold-predicted structures [16] that could be encoded as ESM-3 structure tokens using the EvolutionaryScale esm package, v3.2.1 [3]. For each SAE feature, protein-level maximum activations were extracted using sequence-only (S) and using sequence plus structure (S+St). A paired Wilcoxon signed-rank test (SciPy, v1.13.1, stats.wilcoxon) [31] was performed on the activation distributions, Benjamini-Hochberg FDR correction was applied across all tested features using statsmodels, v0.14.6 [34], and Cohen’s *d_z_* (paired design) was computed as an effect size measure. Features were classified as structure-enhanced (adjusted *p <* 0.05 and *d >* 0.2), structure-suppressed (adjusted *p <* 0.05 and *d <* −0.2), or structure-invariant (all others).

### Centered kernel alignment

How structure tokens reshape ESM-3’s internal representations was measured using linear-centered kernel alignment (CKA) [35], implemented in NumPy v2.1.0 [33]. For 991 proteins with successfully encoded AlphaFold structures [16], residual stream activations were extracted under S-only and S+St conditions at 12 layers spanning the full network (layers 0, 4, 8, 12, 16, 20, 24, 28, 32, 36, 42, 47). CKA was computed between the S and S+St activation matrices at each layer, using only proteins for which both conditions produced valid outputs. A CKA value near 1 indicates that structure tokens leave the representation geometry unchanged; values near 0 indicate that the two conditions produce geometrically dissimilar representations.

### Attention analysis

ESM-3 uses PyTorch’s, v2.6.0+cu124 [29] F.scaled_dot_product_attention, which does not expose attention weights. The weights were extracted by monkey-patching the forward method of each MultiHeadAttention module. To do so, 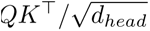 was computed explicitly, softmax was applied, the resulting attention matrices were stored, and the results were passed through the standard value projection. For ESM-2, native HuggingFace transformers, v4.48.1 [30], output_attentions=True flag, were used. Attention matrices were extracted from 494 proteins under both S and S+St conditions for ESM-3 (S-only for ESM-2) in all layers and heads. The structure-responsiveness of each head was quantified as the Jensen-Shannon divergence (JSD) between the attention matrices S and S+St, calculated using SciPy, v1.13.1 [31] and averaged over all proteins and positions; heads with JSD *>* 0.1 were classified as structure-responsive.

For contact prediction, the average product correction (APC) [36] was applied to the symmetrized attention matrices and a supervised logistic regression classifier with penalty *ℓ*_1_ was trained using scikit-learn, v1.6.1 [37] on 500,000 pairs of residues to predict whether two residues are in contact (*C_α_* distance *<* 8 Å).

### Layer and head ablation

The causal importance of individual layers and attention heads was quantified using mean ablation, implemented in PyTorch v2.6.0+cu124 using forward hooks [29]. For layer ablation, a pre-hook was registered in the forward position on the layer *ℓ* + 1 that replaced its input (the output of the layer *ℓ*) with the mean activation calculated over a reference set of 500 proteins. The effect of ablating each layer was measured as the KL divergence between the ablated and unablated output distributions, averaged across all tokens in the reference set. This hook-based design avoids modifying the target layer’s forward pass and works identically for both ESM-2 and ESM-3.

For head ablation, the attention module’s forward method was monkey-patched to zero out the output of a single head before recombining across heads. Because ESM-2 and ESM-3 implement multi-head attention differently (ESM-2 separates heads explicitly; ESM-3 uses fused projections), architecture-specific patches were written that reshape the attention output into (*B, L, H, d_head_*), zero the target head, and reshape back. As with layer ablation, the effect metric was KL divergence from the unablated baseline. All heads in both models were tested (ESM-2: 33 layers × 20 heads = 660; ESM-3: 48 layers × 24 heads = 1,152).

### L0H7 functional impact analysis

To quantify the functional consequences of ablating the geometric attention head L0H7, 100 proteins with AlphaFold-predicted structures were randomly selected (seed = 42) from the evaluation set, filtered to 50–300 residues. Each protein was run through ESM-3 under sequence-plus-structure (S+St) conditions twice: once with the unmodified model and once with L0H7 zeroed using the head ablation procedure described above.

Three functional readouts were compared between the normal and ablated forward passes. First, secondary structure predictions were obtained from ESM-3’s native secondary structure output head (secondary_structure_logits), which produces per-residue logits over three classes (helix, sheet, coil). The top-1 prediction at each position was compared between normal and ablated conditions; the fraction of positions where the predicted class changed was reported as the SS3 agreement metric (1 − agreement = fraction altered). Second, amino acid predictions were compared by taking the top-1 token from ESM-3’s sequence logits (sequence_logits) at each position; the fraction of positions where the predicted amino acid differed between conditions was reported as the token change rate. Third, residual stream activations at layer 33 were extracted under both conditions and encoded through the trained SAE. The mean activation of the top 50 structure-enhanced features (ranked by cross-modal effect size) was computed across all residue positions in each protein; the percentage change in this mean between normal and ablated conditions, averaged across all 100 proteins, yielded the 28% activation drop reported in the text.

To establish specificity, 10 control heads were selected from the remaining 23 layer-0 heads (excluding L0H7) by random sampling without replacement (numpy.random.RandomState, seed = 42). Each control head was ablated individually using the same procedure and protein set, and the same three functional readouts were computed. Results were summarized as the mean ± standard deviation across the 10 control heads.

### Attribution patching

Attribution patching estimates the causal contribution of each layer to a prediction change without requiring a separate forward pass per layer [8]. A set of 200 pairs of proteins was selected, each consisting of one helix-dominant protein and one sheet-dominant protein, with secondary structure assignments calculated using DSSP, v4.5.8 [38]. For each pair, residual stream activations of the helix-dominant protein (source) were cached at each layer using PyTorch, v2.6.0+cu124, forward hooks [29]. A forward pass was then performed on the sheet-dominant protein (target) with gradient hooks enabled. The loss was defined as the KL divergence between the source and target logit distributions, with the source distribution detached from the computation graph. The attribution score for layer *ℓ* was calculated as 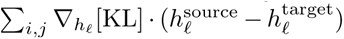, where the sum is distributed over all positions *i* and hidden dimensions *j*. This gradient-based approach computes all layer attributions in a single backward pass.

### Steering vectors

Steering vectors were constructed to test whether SAE features causally mediate model predictions along specific biological axes. For each target property (secondary structure, solvent accessibility), the steering direction was computed as the mean decoder column of the 20 SAE features most enhanced by structure tokens, extracted from the trained SAE weight matrix using PyTorch (v2.6.0+cu124). During inference, a forward hook at layer 33 added *α* × v to the residual stream at every position, where v is the unit-normalized steering direction. Eight strengths were tested: *α* ∊ {−50, −20, −10, −5, 5, 10, 20, 50} on 500 proteins. For each steering condition, KL divergence from the unsteered baseline was measured using PyTorch’s F.kl_div [29], along with the change in activation of the target features, and the fraction of tokens whose top-1 prediction changed.

### Sparse feature circuits

To map causal dependencies between SAE features at different layers [39], a pair-wise ablation procedure was implemented in PyTorch, v2.6.0+cu124 [29]. For each of 50 downstream features in a later layer, the 100 most active upstream features in an earlier layer were identified. Each upstream feature was then individually ablated (by setting its activation to zero before decoding), and the change in downstream feature activation across 20 proteins was measured. A causal edge was retained if the change was significant using a Bonferroni-corrected *t*-test (SciPy, v1.13.1, stats.ttest_1samp) [31] *and* the effect size exceeded the median plus two standard deviations of the null distribution. Three-layer pairs were tested: ESM-3 layer 16→33 and 33→42, and ESM-2 layer 16→24.

### Negative controls

Three negative controls were implemented to test whether SAE features carry genuine biological information beyond what simpler decompositions provide. *Random projection*: sparse random features matching the SAE’s sparsity (*k* = 64 non-zero entries per token), with magnitudes sampled from the empirical distribution of SAE activations, generated using NumPy, v2.1.0 [33], with fixed random seeds. *Shuffled features*: column-wise permutation of residue indices within each feature using NumPy’s random permutation, preserving per-feature activation statistics but destroying residue-level correspondence. A PCA baseline was established from the top 64 principal components of the raw activation matrix, which was computed using scikit-learn, v1.6.1 (decomposition.PCA) [37]. Random projection and shuffled baselines were repeated with 3 random seeds. All comparisons used linear classifiers (scikit-learn LogisticRegression with *ℓ*_2_ penalty) with splits at protein-level train/test (proteins, not residues, were randomly assigned) to prevent leakage of correlated residues within a single protein.

### Bootstrap confidence intervals

All reported confidence intervals were calculated from 2,000 bootstrap iterations with protein-level resampling using NumPy, v2.1.0 [33]. Proteins were drawn with replacement, and all residues belonging to each sampled protein were included. The intervals are bias-corrected and accelerated (BCa) [40], calculated using SciPy, v1.13.1, stats.bootstrap [31], where the jackknife influence function was computationally tractable and percentile intervals otherwise. All intervals are reported at the 95% confidence level.

### Software and hardware

All analyses were performed in Python v3.11 (https://www.python.org/) with PyTorch [29], scikit-learn [37], SciPy [31], and gokit (https://github.com/JLSteenwyk/gokit) for GO enrichment. GPU computations were performed on two NVIDIA RTX 6000 Ada GPUs (48 GB each) on a workstation with an AMD Threadripper PRO 7995WX 96-core processor and 1 TB of RAM. ESM-3 forward passes were distributed across both GPUs for proteins with more than 512 residues; all other computations used a single GPU. Figures are made with colorblind-friendly color palettes using pypubfigs, v1.1.3, a Python implementation of ggpubfigs [41].

## Data and code availability

All data and code supporting this study are available at Figshare (DOI: 10.6084/m9.figshare.32059860; the following link is provided for review purposes only: https://figshare.com/s/7cfd5d831dde153cef28). The repository contains: (i) 1.5 million UniRef50 training sequences and 12,491 Swiss-Prot evaluation proteins with annotations; (ii) 14 trained SAE checkpoints that cover expansion factors 2-8 and *k* = 32, 64 at residue and protein granularity for ESM-2 and ESM-3; (iii) all analysis results files reported in the manuscript; (iv) SAE protein-level activation summaries for both models; and (v) the complete analysis and figure generation pipeline. Intermediate files that can be regenerated from the sequences and models provided (raw ESM activations, cross-modal activations, AlphaFold structures, and unified pipeline intermediates; ∼39 GB total) are not included but can be produced using the scripts provided, as documented in the repository README.

## Extended Data

**Extended Data Fig. 1.**
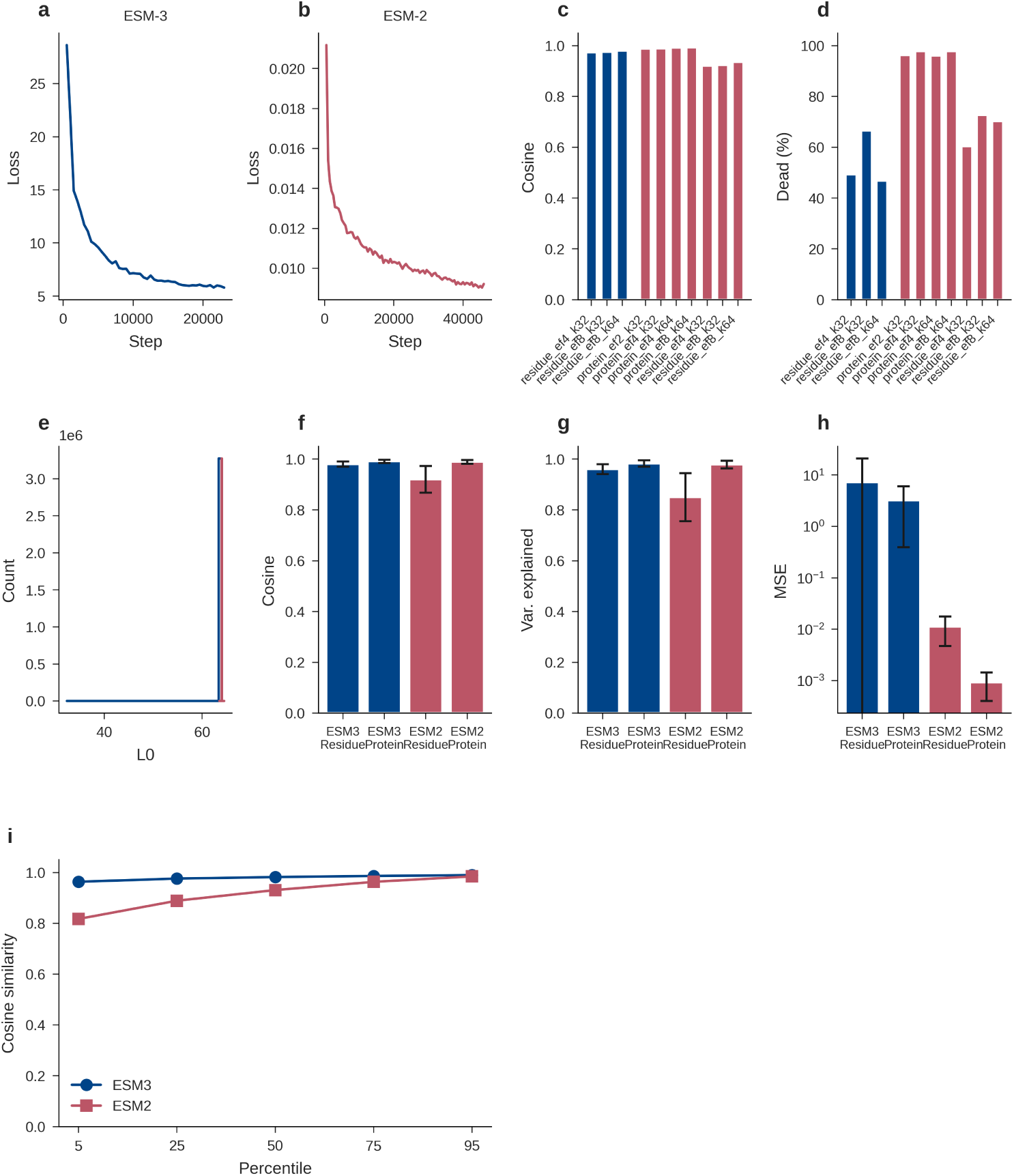
SAE training diagnostics and reconstruction quality. **a,b.** Training loss curves for ESM-3 (**a**) and ESM-2 (**b**) SAEs. Separate y-axes reflect a three-order-of-magnitude difference in absolute loss, arising from ESM-2’s smaller activation dimensionality (*d* = 1,280 compared to 1,536). Both models converge. **c.** Reconstruction cosine similarity across SAE configurations (expansion factors 4 and 8, *k* = 32 and 64, residue- and protein-level). Reconstruction quality is robust to hyperparameter choice, with all configurations achieving cosine similarity above 0.9. This observation indicates that SAE reconstruction quality is robust to hyperparameter choice. **d.** Dead neuron fraction per configuration. ESM-3 SAEs have higher dead fractions than ESM-2, consistent with its larger native dimensionality. **e.** *L*_0_ sparsity histogram confirming TopK enforcement yields exactly *k* = 64 active features per input token. **f.** Reconstruction cosine similarity at residue and protein levels for both models with standard deviation error bars. **g.** Fraction of activation variance explained by SAE reconstructions. **h.** Examination of mean squared error (log *y*-axis) values reveals that ESM-2 residue-level MSE value is lower than ESM-3, reflecting the smaller activation space. **i.** Cosine similarity across percentiles (5th-95th). ESM-3 maintains high similarity even at the 5th percentile (0.96), while ESM-2’s lower tail extends to 0.81.

**Extended Data Fig. 2.**
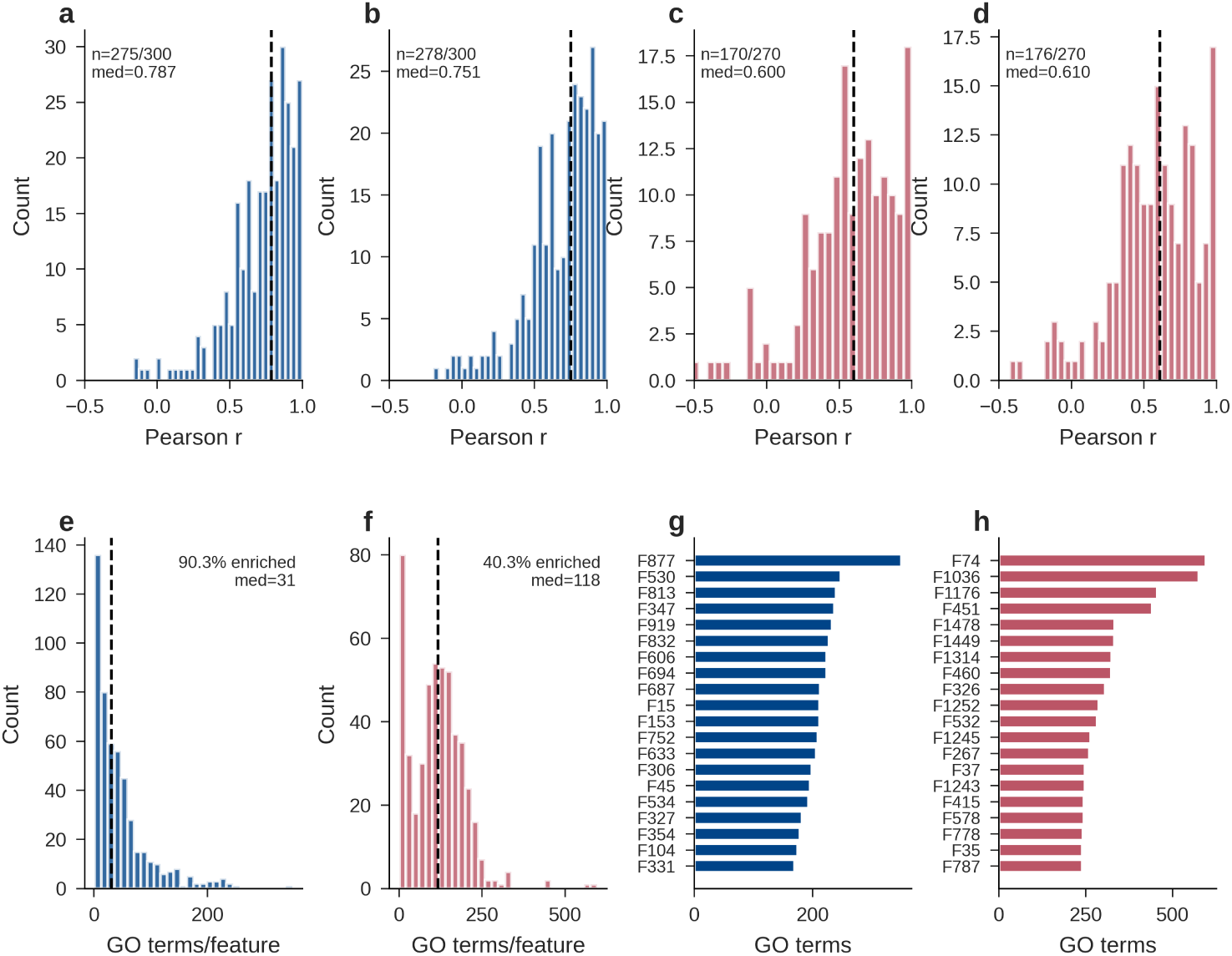
Autointerpretability validation and GO enrichment detail. a-d. Histograms of Pearson *r* between LLM-predicted and observed feature activations on held-out proteins for four judge*×*model conditions: ESM-3/Claude Sonnet 4.6 (median *r* = 0.79, *n* = 275*/*300; **a**), ESM-3/GPT-5.4 (median *r* = 0.75, *n* = 278*/*300; **b**), ESM-2/Claude (median *r* = 0.60, *n* = 170*/*270; **c**), and ESM-2/GPT (median *r* = 0.61, *n* = 176*/*270; **d**). Dashed lines mark medians. The *n/N* counts reflect features for which the LLM returned parseable predictions; the remaining *N − n* failed to produce valid JSON and were excluded. ESM-3 features are consistently more interpretable than ESM-2 features across both judges. **e,f.** Distribution of the number of significantly enriched GO terms per feature (BH-FDR *<* 0.05) for ESM-3 (**e**; 90.3% of features enriched, median 31 terms among enriched features) and ESM-2 (**f**; 40.3% enriched, median 118 terms). ESM-2’s higher median among enriched features reflects the fact that only its most broadly activating features pass the enrichment threshold; the majority are too sparse for the test to detect enrichment. **g,h.** Top 20 features ranked by number of enriched GO terms for ESM-3 (**g**) and ESM-2 (**h**). The most GO-enriched individual features carry hundreds of functional annotations.

**Extended Data Fig. 3.**
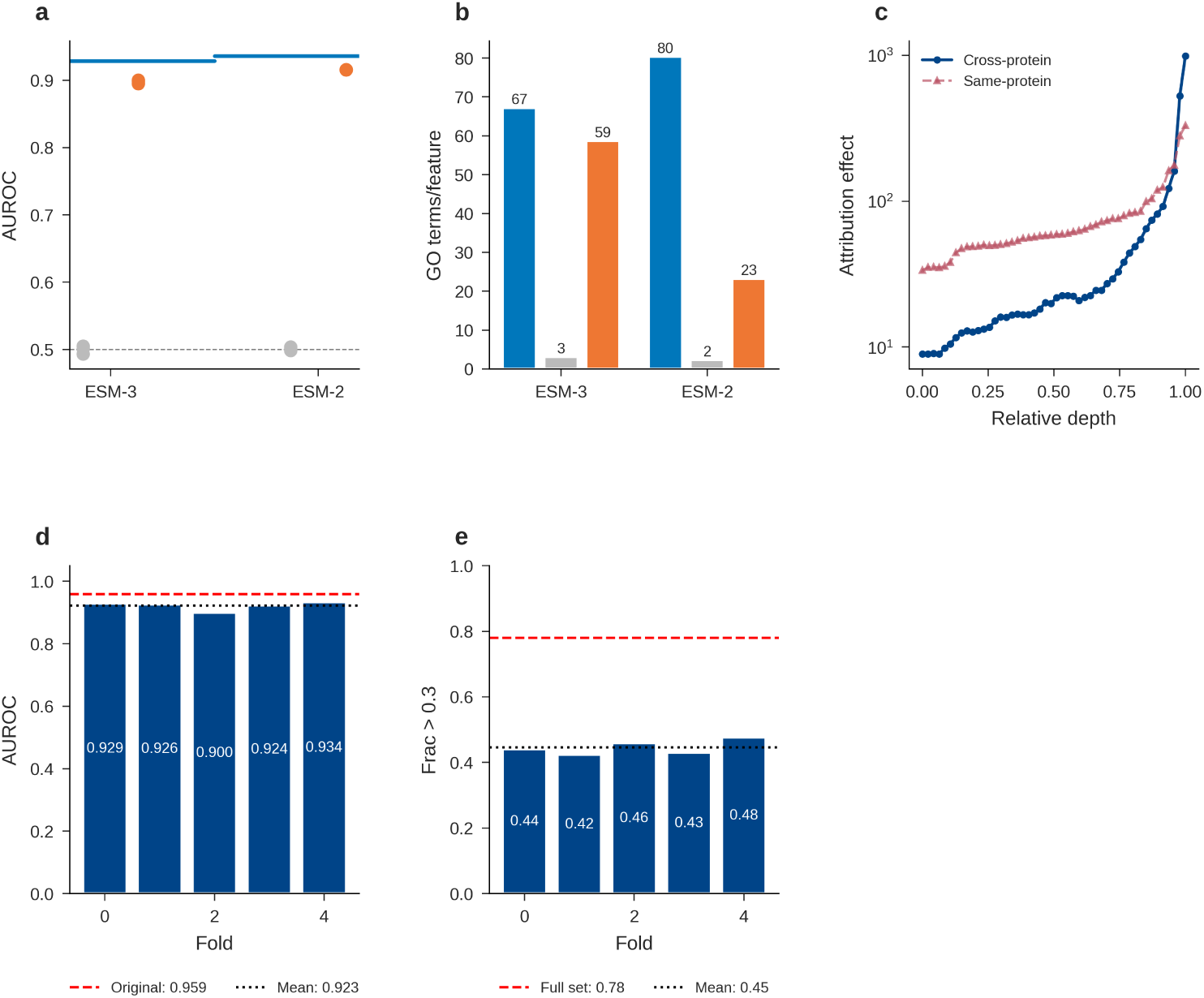
Negative controls and robustness analyses. **a.** Functional site detection AUROC for learned SAE features (blue horizontal lines) compared with random-direction projections (grey circles) and label-shuffled baselines (orange circles) for both ESM-3 and ESM-2. SAE features (0.93-0.94) far exceed random projections (0.50), confirming that learned features carry genuine biological information. Dashed line at 0.5 (chance level). **b.** Mean number of significantly enriched GO terms per feature for SAE features (blue) compared with random projection (orange) and shuffled (grey) baselines. SAE features yield 20-30*×* more enriched GO terms than controls. **c.** Attribution patching effect sizes for cross-protein (blue, solid) versus same-protein S/S+St (red, dashed) interventions as a function of relative network depth. Both show similar late-layer peaks, confirming that the attribution profile reflects structure processing rather than sequence differences between paired proteins. **d.** Probing AUROC under a homology-aware 50% sequence identity split across five folds. Mean fold AUROC (0.923, black dotted line) is close to the original single-split AUROC (0.959, red dashed line), confirming that results are not driven by sequence similarity between training and test proteins. **e.** Fraction of convergent features (*r >* 0.3) under the same homology split. Mean across folds (0.45, black dotted line) compared with the full dataset (0.78, red dashed line). The lower fraction reflects the more stringent split removing homologous proteins that inflate convergence estimates.

**Extended Data Fig. 4.**
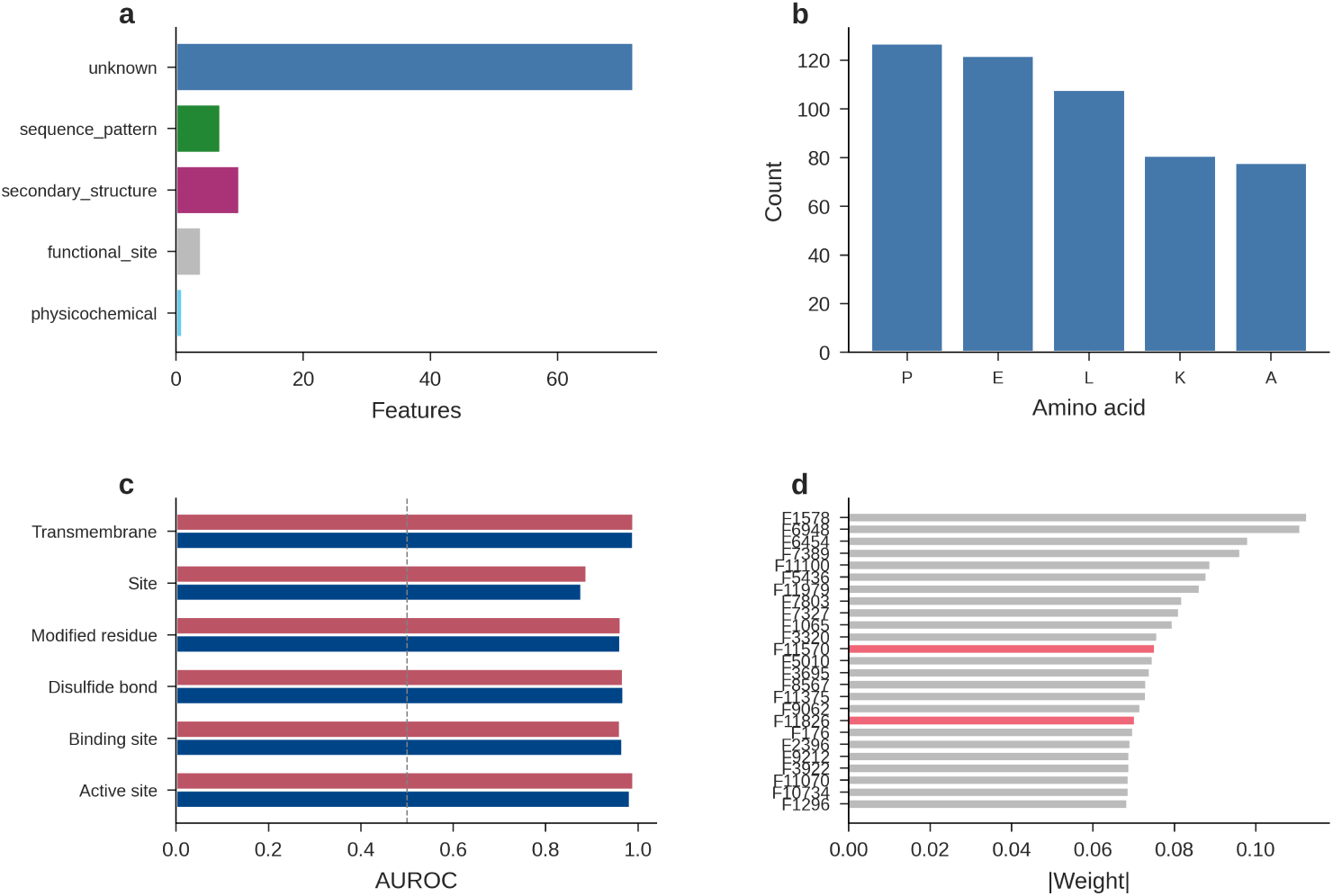
Feature biology and residue-level characterisation. **a.** Distribution of annotated motif types across SAE features classified by the autointerpretability pipeline. The majority of features are classified as unknown, reflecting the conservative nature of the automated annotation; the remainder span secondary structure, sequence pattern, functional site, and physico-chemical categories. **b.** Amino acid composition of a representative SAE feature, showing the top 5 most frequently activated residue types. This feature preferentially activates on proline (P), glutamate (E), and leucine (L), illustrating residue-type selectivity of individual SAE features. **c.** Per-residue-type functional site detection AUROC for ESM-3 (blue) and ESM-2 (red) across six annotation categories: active site, binding site, disulfide bond, modified residue, site, and transmembrane. Both models achieve AUROC above 0.85 for all categories, with ESM-3 slightly outperforming ESM-2 on most types. Dashed line at 0.5 (chance). **d.** Top 25 SAE features ranked by absolute linear probe weight (*|w|*), coloured by cross-modal category: structure-enhanced (red), structure-suppressed (blue), and invariant (grey). Structure-responsive features are interspersed among the top probe features, indicating that both structure-sensitive and structure-invariant features contribute to functional site detection.

**Extended Data Fig. 5.**
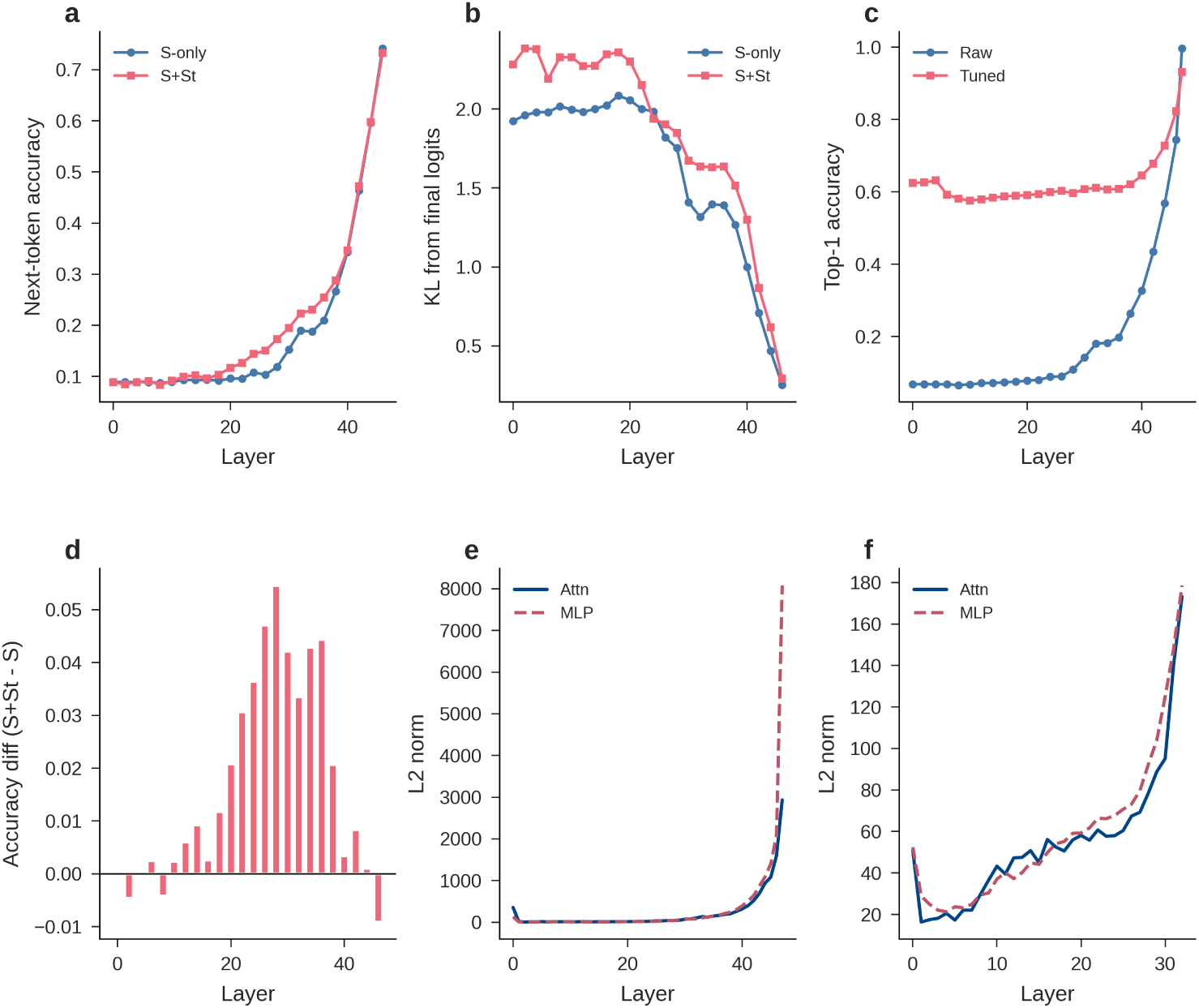
Logit lens and residual stream decomposition. **a.** Logit lens next-token prediction accuracy across ESM-3 layers under sequence-only (S, blue) and sequence-plus-structure (S+St, red) conditions. Accuracy rises steeply after layer 30, with S+St consistently outperforming S-only, indicating that structure tokens improve prediction formation in late layers. **b.** KL divergence between logit lens predictions and the final-layer output distribution across ESM-3 layers. KL decreases sharply in late layers as intermediate representations approach the final prediction. S+St shows lower KL than S-only in the final layers, consistent with structure tokens aiding prediction formation. **c.** Comparison of raw logit lens (blue) and tuned lens (red) top-1 accuracy across ESM-3 layers. The tuned lens, which applies a learned affine transformation at each layer, substantially outperforms the raw logit lens, particularly in mid-layers where representations have not yet aligned with the output vocabulary. **d.** Per-layer accuracy difference (S+St minus S) for the logit lens. Structure tokens provide the largest accuracy gains in layers 20-40, coinciding with the integration zone identified by CKA analysis (Fig. 3a). **e.** *L*_2_ norms of attention (solid) and MLP (dashed) output contributions to the residual stream at each layer for ESM-3. MLP contributions spike dramatically in the final layer, indicating that the output projection relies heavily on MLP computation. **f.** Same analysis for ESM-2. Both attention and MLP contributions increase through the network, with MLP dominating in late layers. The scale is two orders of magnitude smaller than ESM-3, reflecting ESM-2’s smaller model dimensionality.

**Extended Data Fig. 6.**
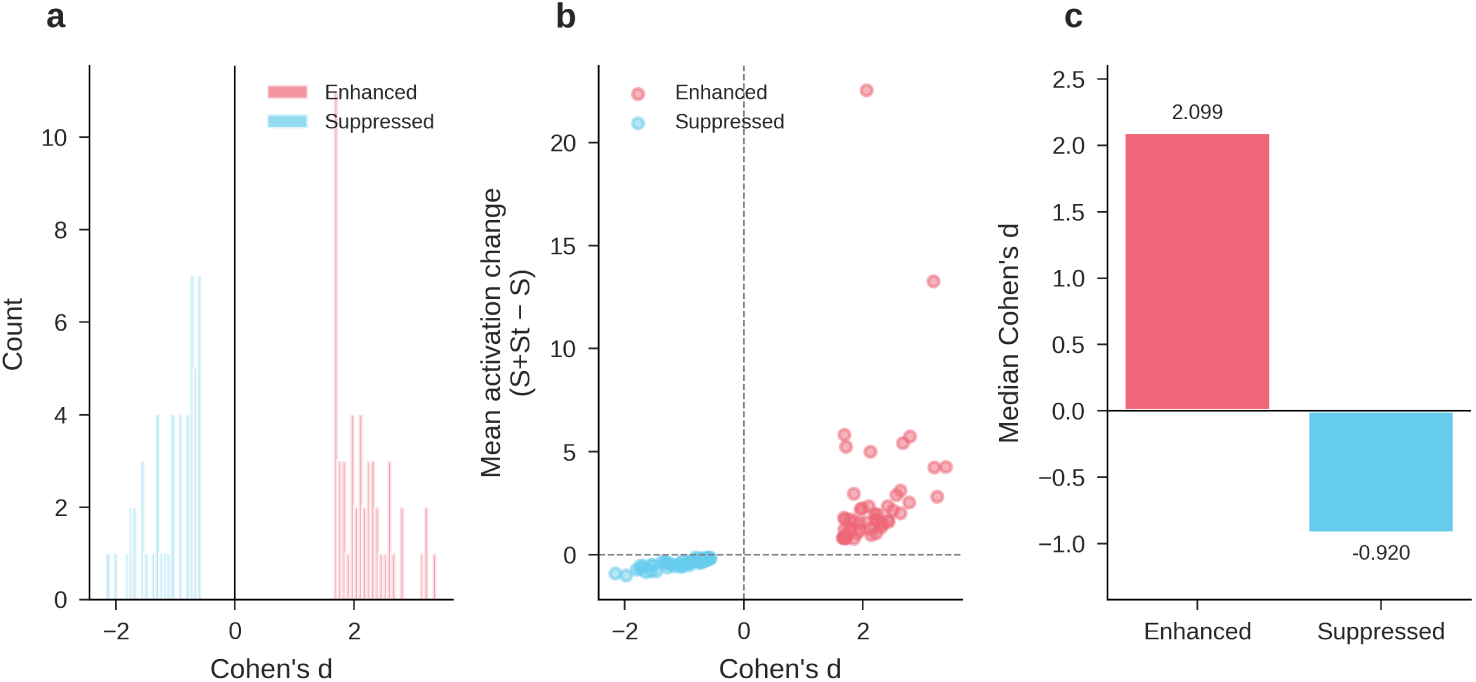
Cross-modal feature response detail. **a.** Distribution of Cohen’s *d* effect sizes for the top 50 structure-enhanced (red) and top 50 structure-suppressed (blue) SAE features. Enhanced features cluster at positive *d* values (median 2.10), while suppressed features cluster at negative values (median *−*0.92). The separation confirms that the cross-modal classification reflects genuine differences in activation magnitude, not statistical noise. **b.** Cohen’s *d* compared to mean activation change (S+St minus S) for each feature. Enhanced features (red) show positive activation changes that scale with effect size, while suppressed features (blue) cluster near zero on both axes. The relationship is monotonic, indicating that Cohen’s *d* captures biologically meaningful variation in structure responsiveness. **c.** Median Cohen’s *d* by cross-modal category. Enhanced features have a median *d* of 2.10; suppressed features have a median *d* of *−*0.92. The larger absolute effect size for enhanced features indicates that structure tokens produce stronger amplification than suppression of SAE feature activations.

**Extended Data Fig. 7.**
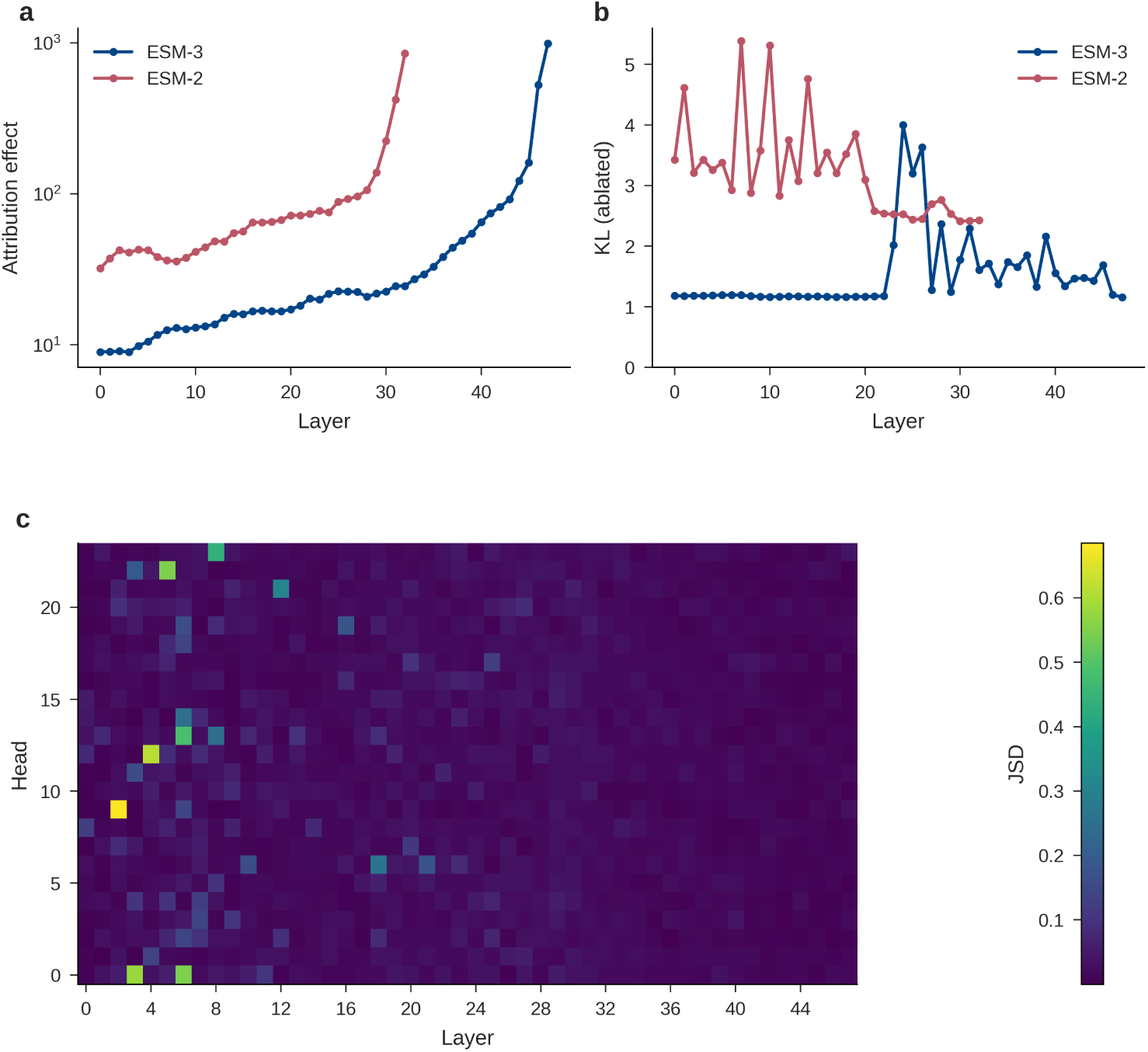
Attribution patching, layer ablation, and attention atlas. **a.** Attribution patching effect size by absolute layer position for ESM-3 (blue) and ESM-2 (red) on a log *y*-axis. Both models concentrate causal contributions in their final layers, with effects increasing by two orders of magnitude from early to late layers. ESM-2 starts with a higher baseline effect, consistent with the absence of a structure embedding that would offload early-layer computation. **b.** KL divergence from mean-ablating each layer individually for ESM-3 (blue) and ESM-2 (red). ESM-3 shows concentrated vulnerability at layers 24-26 (KL up to 4.0), immediately preceding the integration zone, while ESM-2 exhibits more distributed sensitivity with peaks at layers 1, 7, and 14. The ESM-3 peak aligns with the representational phase transition identified by within-model CKA (Fig. 3b). **c.** Full attention head atlas (48 layers *×* 24 heads) showing Jensen-Shannon divergence between S and S+St attention patterns, averaged across 494 proteins. Twenty-eight structure-responsive heads (JSD *>* 0.1) cluster in layers 2-9, with the brightest signal at L0H7 (JSD = 0.71). No responsive heads appear in layers 32-47, confirming that attention-level structure responsiveness is confined to the early network while late layers process structure information through the residual stream rather than through attention pattern changes.

**Extended Data Fig. 8.**
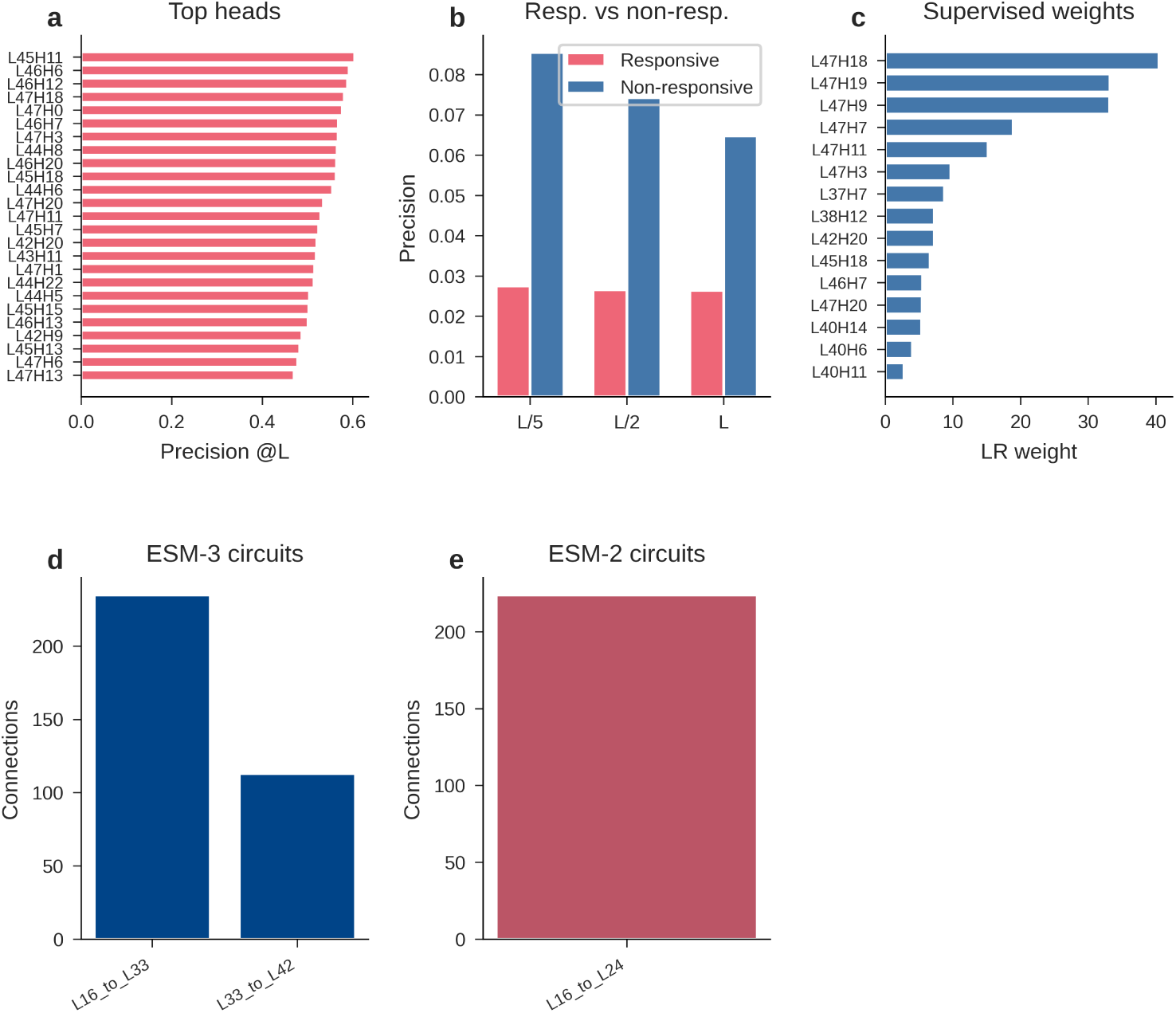
Contact prediction from attention and sparse feature circuits. **a.** Top 25 contact-predicting attention heads ranked by precision at sequence separation *|i − j| ≥ L* (full protein length). Late-layer heads dominate, with L45H11 achieving the highest precision (0.60). All top heads reside in layers 42-47, indicating that contact prediction is concentrated in the final network layers. **b.** Contact prediction precision at three sequence separation thresholds (*L/*5, *L/*2, *L*) for structure-responsive heads (red) compared to non-responsive heads (blue). Non-responsive heads outperform responsive heads at all thresholds, confirming that the heads that read structure in (layers 2-9) are distinct from those that predict contacts out (layers 44-47). **c.** Supervised logistic regression weights assigned to individual attention heads for contact prediction. The highest-weighted heads (L47H18, L47H19, L47H9) are all in the final layers, consistent with the precision ranking in **a**. **d.** Sparse feature circuit connections for ESM-3. The L16*→*L33 layer pair yields 235 significant causal connections, while L33*→*L42 yields 113, reflecting decreasing connection density as representations converge in later layers. **e.** Sparse feature circuit connections for ESM-2. The L16*→*L24 layer pair yields 224 connections, comparable to ESM-3’s early-to-mid density, suggesting similar causal wiring across architectures.

**Extended Data Fig. 9.**
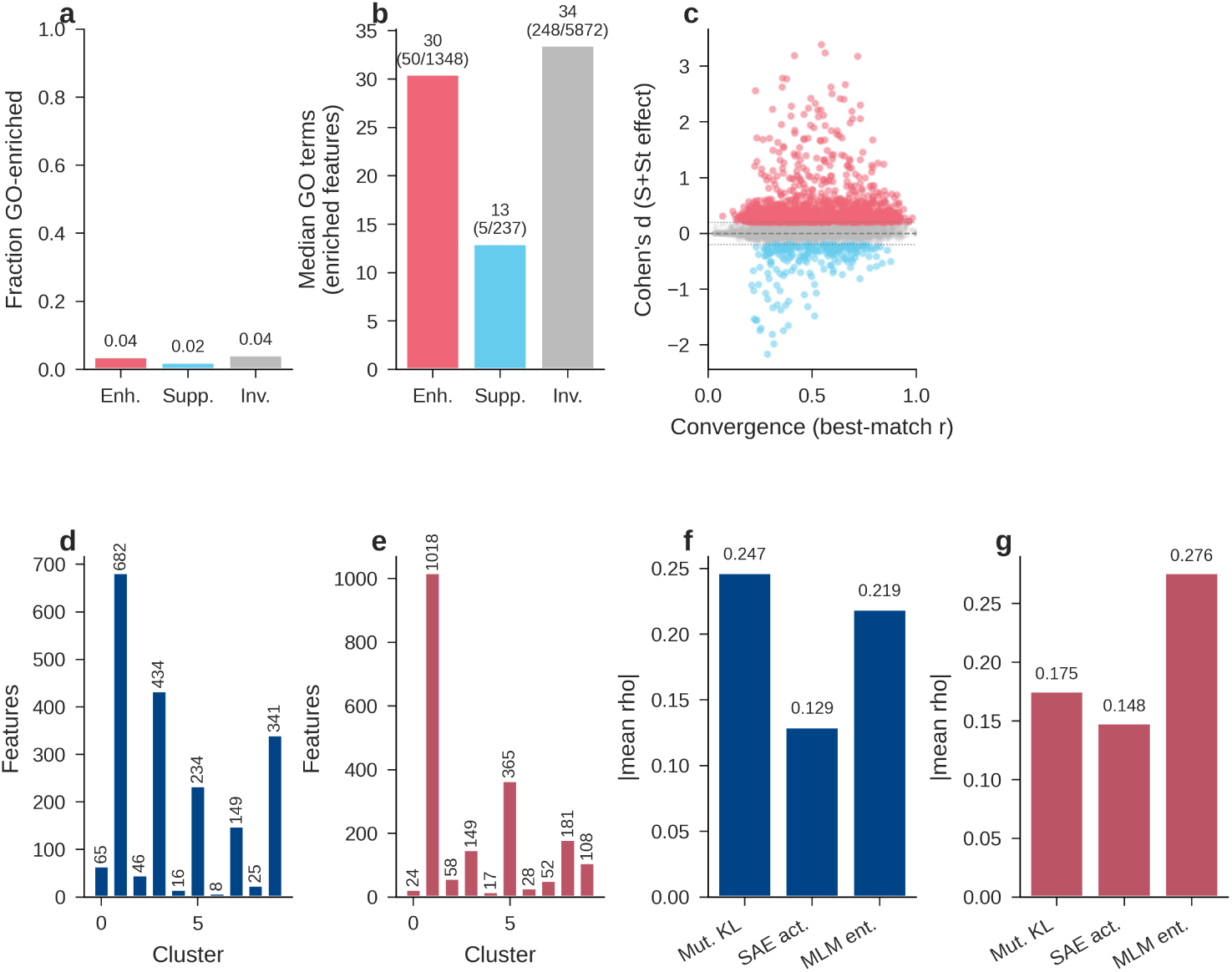
Feature-property relationships, coactivation, and deep mutational scanning. **a.** Fraction of GO-enriched features by cross-modal category: enhanced (4%), suppressed (2%), and invariant (4%). The low fractions reflect that GO enrichment was computed for a subset of features; the comparable rates across categories indicate that GO enrichment is not concentrated in any one category. **b.** Median number of enriched GO terms among GO-enriched features by category, with the number of enriched features out of total features annotated above each bar. Enhanced: 30 terms (50/1,348); suppressed: 13 (5/237); invariant: 34 (248/5,872). Despite comparable enrichment rates, invariant features carry slightly more GO terms per feature, while suppressed features carry the fewest. **c.** Cohen’s *d* (cross-modal effect size) compared to convergence score (best-match *r* with ESM-2) for each feature, colored by cross-modal category: enhanced (red), suppressed (blue), invariant (grey). Enhanced features cluster at higher convergence values, visualizing the paradox that structure-responsive features are more similar to sequence-only ESM-2. Dashed lines at *d* = *±*0.2 mark the classification threshold. **d,e.** Coactivation cluster size distributions (*k* = 10 nearest-neighbor clustering) for ESM-3 (**d**) and ESM-2 (**e**). Both models show a dominant cluster containing most features, along with several smaller, specialized clusters. **f,g.** Correlation between deep mutational scanning fitness effects and three interpretability metrics — mutation-induced KL divergence, SAE feature activation change, and masked language model entropy — for ESM-3 (**f**) and ESM-2 (**g**). Values are mean absolute Spearman *|ρ|* across DMS datasets. Mutation KL and MLM entropy show stronger correlations than SAE activation change, suggesting that global model perturbation metrics capture fitness effects more directly than individual feature activations.

**Extended Data Fig. 10.**
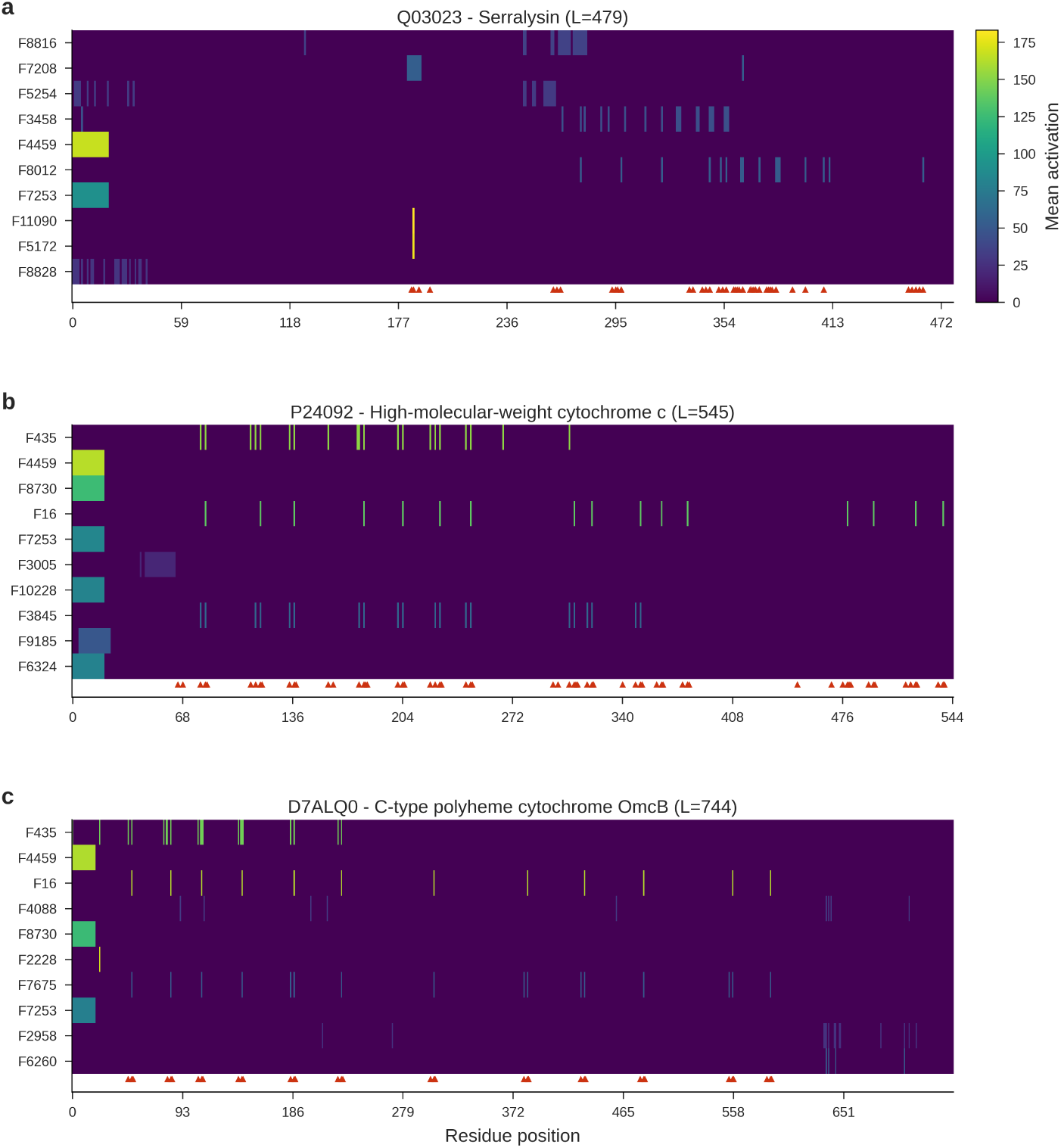
SAE features activate at annotated functional sites. Each panel shows per-residue SAE feature activation profiles for the top 10 features (rows) across the full protein sequence (columns). Red triangles below each heatmap mark experimentally annotated functional sites (active sites, binding sites, metal binding, modified residues) from UniProt. Viridis colormap; shared colorbar scaled to panel **a**. **a.** Q03023, serralysin from *Pseudomonas aeruginosa* (479 aa; 42 active site and binding site residues). Feature activations align with the red triangles at the zinc-binding site (position 177) and the calcium-coordinating residues (positions 295–413), demonstrating that structure-enhanced features (F4459, F7253) localize to residues whose function depends on three-dimensional positioning. **b.** P24092, high-molecular-weight cytochrome *c* from *Desulfovibrio vulgaris* (545 aa; 63 binding sites). Features F435 and F16 show periodic activations that track the regularly spaced heme-binding CXXCH motifs characteristic of multi-heme cytochromes. Red triangles confirm these activations coincide with annotated binding residues. **c.** D7ALQ0, C-type polyheme cytochrome OmcB from *Geobacter sulfurreducens* (744 aa; 37 binding and lipidation sites). The same periodic heme-binding pattern is captured by the same shared features (F435, F16), indicating that the SAE has learned a general heme-coordination feature that generalizes across cytochrome families.

## References

[1] Lin, Z., Akin, H., Rao, R., Hie, B., Zhu, Z., Lu, W., Smetanin, N., Verkuil, R., Kabeli, O., Shmueli, Y., et al.: Evolutionary-scale prediction of atomic-level protein structure with a language model. Science 379(6637), 1123–1130 (2023)

[2] Rives, A., Meier, J., Sercu, T., Goyal, S., Lin, Z., Liu, J., Guo, D., Ott, M., Zitnick, C.L., Ma, J., et al.: Biological structure and function emerge from scaling unsupervised learning to 250 million protein sequences. Proceedings of the national academy of sciences 118(15), 2016239118 (2021)

[3] Hayes, T., Rao, R., Akin, H., Sofroniew, N.J., Oktay, D., Lin, Z., Verkuil, R., Tran, V.Q., Deaton, J., Wiggert, M., et al.: Simulating 500 million years of evolution with a language model. Science 387(6736), 850–858 (2025)

[4] Meier, J., Rao, R., Verkuil, R., Liu, J., Sercu, T., Rives, A.: Language models enable zero-shot prediction of the effects of mutations on protein function. Advances in neural information processing systems 34, 29287–29303 (2021)

[5] Nijkamp, E., Ruffolo, J.A., Weinstein, E.N., Naik, N., Madani, A.: Progen2: exploring the boundaries of protein language models. Cell systems 14(11), 968–978 (2023)

[6] Olah, C., Cammarata, N., Schubert, L., Goh, G., Petrov, M., Carter, S.: Zoom in: An introduction to circuits. Distill 5(3), 00024–001 (2020)

[7] Elhage, N., Hume, T., Olsson, C., Schiefer, N., Henighan, T., Kravec, S., Hatfield-Dodds, Z., Lasenby, R., Drain, D., Chen, C., et al.: Toy models of superposition. arXiv preprint arXiv:2209.10652 (2022)

[8] Nanda, N., Chan, L., Lieberum, T., Smith, J., Steinhardt, J.: Progress measures for grokking via mechanistic interpretability. arXiv preprint arXiv:2301.05217 (2023)

[9] Simon, E., Zou, J.: Interplm: discovering interpretable features in protein language models via sparse autoencoders. Nature methods 22(10), 2107–2117 (2025)

[10] Gujral, O., Bafna, M., Alm, E., Berger, B.: Sparse autoencoders uncover biologically interpretable features in protein language model representations. Proceedings of the National Academy of Sciences 122(34), 2506316122 (2025)

[11] Cunningham, H., Ewart, A., Riggs, L., Huben, R., Sharkey, L.: Sparse autoencoders find highly interpretable features in language models. arXiv preprint arXiv:2309.08600 (2023)

[12] Templeton, A., Conerly, T., Marcus, J., Lindsey, J., Bricken, T., Chen, B., Pearce, A., Citro, C., Ameisen, E., Jones, A., Cunningham, H., Turner, N.L., McDougall, C., MacDiarmid, M., Freeman, C.D., Sumers, T.R., Rees, E., Batson, J., Jermyn, A., Carter, S., Olah, C., Henighan, T.: Scaling monosemanticity: Extracting interpretable features from claude 3 sonnet. Transformer Circuits Thread (2024)

[13] Bricken, T., Templeton, A., Batson, J., Chen, B., Jermyn, A., Conerly, T., Turner, N., Anil, C., Denison, C., Askell, A., et al.: Towards monosemanticity: Decomposing language models with dictionary learning. Transformer Circuits Thread 2(5), 6 (2023)

[14] Rao, R., Bhattacharya, N., Thomas, N., Duan, Y., Chen, P., Canny, J., Abbeel, P., Song, Y.: Evaluating protein transfer learning with tape. Advances in neural information processing systems 32 (2019)

[15] Vig, J., Madani, A., Varshney, L.R., Xiong, C., Socher, R., Rajani, N.F.: Bertology meets biology: Interpreting attention in protein language models. arXiv preprint arXiv:2006.15222 (2020)

[16] Jumper, J., Evans, R., Pritzel, A., Green, T., Figurnov, M., Ronneberger, O., Tunyasuvunakool, K., Bates, R., Žídek, A., Potapenko, A., et al.: Highly accurate protein structure prediction with alphafold. nature 596(7873), 583–589 (2021)

[17] Ding, F., Denain, J.-S., Steinhardt, J.: Grounding representation similarity through statistical testing. Advances in Neural Information Processing Systems 34, 1556–1568 (2021)

[18] Ferruz, N., Schmidt, S., Höcker, B.: Protgpt2 is a deep unsupervised language model for protein design. Nature communications 13(1), 4348 (2022)

[19] Su, J., Han, C., Zhou, Y., Shan, J., Zhou, X., Yuan, F.: Saprot: Protein language modeling with structure-aware vocabulary. BioRxiv, 2023–10 (2023)

[20] Blum, M., Andreeva, A., Florentino, L.C., Chuguransky, S.R., Grego, T., Hobbs, E., Pinto, B.L., Orr, A., Paysan-Lafosse, T., Ponamareva, I., et al.: Interpro: the protein sequence classification resource in 2025<? mode longmeta?>. Nucleic acids research 53(D1), 444–456 (2025)

[21] Su, J., Ahmed, M., Lu, Y., Pan, S., Bo, W., Liu, Y.: Roformer: Enhanced transformer with rotary position embedding. Neurocomputing 568, 127063 (2024)

[22] Suzek, B.E., Wang, Y., Huang, H., McGarvey, P.B., Wu, C.H., UniProt Consortium, t.: Uniref clusters: a comprehensive and scalable alternative for improving sequence similarity searches. Bioinformatics 31(6), 926–932 (2015)

[23] Boeckmann, B., Bairoch, A., Apweiler, R., Blatter, M.-C., Estreicher, A., Gasteiger, E., Martin, M.J., Michoud, K., O’Donovan, C., Phan, I., et al.: The swiss-prot protein knowledgebase and its supplement trembl in 2003. Nucleic acids research 31(1), 365–370 (2003)

[24] Steinegger, M., Söding, J.: Mmseqs2 enables sensitive protein sequence searching for the analysis of massive data sets. Nature biotechnology 35(11), 1026–1028 (2017)

[25] Aleksander, S.A., Balhoff, J., Carbon, S., Cherry, J.M., Drabkin, H.J., Ebert, D., Feuermann, M., Gaudet, P., Harris, N.L., et al.: The gene ontology knowledgebase in 2023. Genetics 224(1), 031 (2023)

[26] Mistry, J., Chuguransky, S., Williams, L., Qureshi, M., Salazar, G.A., Sonnhammer, E.L., Tosatto, S.C., Paladin, L., Raj, S., Richardson, L.J., et al.: Pfam: The protein families database in 2021. Nucleic acids research 49(D1), 412–419 (2021)

[27] Uniprot: the universal protein knowledgebase in 2021. Nucleic acids research 49(D1), 480–489 (2021)

[28] Gao, L., Tour, T.D., Tillman, H., Goh, G., Troll, R., Radford, A., Sutskever, I., Leike, J., Wu, J.: Scaling and evaluating sparse autoencoders. arXiv preprint arXiv:2406.04093 (2024)

[29] Paszke, A., Gross, S., Massa, F., Lerer, A., Bradbury, J., Chanan, G., Killeen, T., Lin, Z., Gimelshein, N., Antiga, L., et al.: Pytorch: An imperative style, high-performance deep learning library. Advances in neural information processing systems 32 (2019)

[30] Wolf, T., Debut, L., Sanh, V., Chaumond, J., Delangue, C., Moi, A., Cistac, P., Rault, T., Louf, R., Funtowicz, M., et al.: Transformers: State-of-the-art natural language processing. In: Proceedings of the 2020 Conference on Empirical Methods in Natural Language Processing: System Demonstrations, pp. 38–45 (2020)

[31] Virtanen, P., Gommers, R., Oliphant, T.E., Haberland, M., Reddy, T., Cournapeau, D., Burovski, E., Peterson, P., Weckesser, W., Bright, J., et al.: Scipy 1.0: fundamental algorithms for scientific computing in python. Nature methods 17(3), 261–272 (2020)

[32] Benjamini, Y., Hochberg, Y.: Controlling the false discovery rate: a practical and powerful approach to multiple testing. Journal of the Royal statistical society: series B (Methodological) 57(1), 289–300 (1995)

[33] Harris, C.R., Millman, K.J., Van Der Walt, S.J., Gommers, R., Virtanen, P., Cournapeau, D., Wieser, E., Taylor, J., Berg, S., Smith, N.J., et al.: Array programming with numpy. nature 585(7825), 357–362 (2020)

[34] Seabold, S., Perktold, J., et al.: Statsmodels: econometric and statistical modeling with python. scipy 7(1), 92–96 (2010)

[35] Kornblith, S., Norouzi, M., Lee, H., Hinton, G.: Similarity of neural network representations revisited. In: International Conference on Machine Learning, pp. 3519–3529 (2019). PMlR

[36] Dunn, S.D., Wahl, L.M., Gloor, G.B.: Mutual information without the influence of phylogeny or entropy dramatically improves residue contact prediction. Bioinformatics 24(3), 333–340 (2008)

[37] Pedregosa, F., Varoquaux, G., Gramfort, A., Michel, V., Thirion, B., Grisel, O., Blondel, M., Prettenhofer, P., Weiss, R., Dubourg, V., et al.: Scikit-learn: Machine learning in python. the Journal of machine Learning research 12, 2825–2830 (2011)

[38] Kabsch, W., Sander, C.: Dictionary of protein secondary structure: pattern recognition of hydrogen-bonded and geometrical features. Biopolymers: Original Research on Biomolecules 22(12), 2577–2637 (1983)

[39] Marks, S., Rager, C., Michaud, E.J., Belinkov, Y., Bau, D., Mueller, A.: Sparse feature circuits: Discovering and editing interpretable causal graphs in language models. arXiv preprint arXiv:2403.19647 (2024)

[40] Efron, B.: Better bootstrap confidence intervals. Journal of the American statistical Association 82(397), 171–185 (1987)

[41] Steenwyk, J.L., Rokas, A.: ggpubfigs: colorblind-friendly color palettes and ggplot2 graphic system extensions for publication-quality scientific figures. Microbiology Resource Announcements 10(44), 10–1128 (2021)

